# Multimodal subspace identification for modeling discrete-continuous spiking and field potential population activity

**DOI:** 10.1101/2023.05.26.542509

**Authors:** Parima Ahmadipour, Omid G. Sani, Bijan Pesaran, Maryam M. Shanechi

## Abstract

Learning dynamical latent state models for multimodal spiking and field potential activity can reveal their collective low-dimensional dynamics and enable better decoding of behavior through multimodal fusion. Toward this goal, developing unsupervised learning methods that are computationally efficient is important, especially for real-time learning applications such as brain-machine interfaces (BMIs). However, efficient learning remains elusive for multimodal spike-field data due to their heterogeneous discrete-continuous distributions and different timescales. Here, we develop a multiscale subspace identification (multiscale SID) algorithm that enables computationally efficient modeling and dimensionality reduction for multimodal discrete-continuous spike-field data. We describe the spike-field activity as combined Poisson and Gaussian observations, for which we derive a new analytical subspace identification method. Importantly, we also introduce a novel constrained optimization approach to learn valid noise statistics, which is critical for multimodal statistical inference of the latent state, neural activity, and behavior. We validate the method using numerical simulations and spike-LFP population activity recorded during a naturalistic reach and grasp behavior. We find that multiscale SID accurately learned dynamical models of spike-field signals and extracted low-dimensional dynamics from these multimodal signals. Further, it fused multimodal information, thus better identifying the dynamical modes and predicting behavior compared to using a single modality. Finally, compared to existing multiscale expectation-maximization learning for Poisson-Gaussian observations, multiscale SID had a much lower computational cost while being better in identifying the dynamical modes and having a better or similar accuracy in predicting neural activity. Overall, multiscale SID is an accurate learning method that is particularly beneficial when efficient learning is of interest.

## 1. Introduction

Studies of neural population dynamics have mostly focused on a single modality of neural activity such as spikes or field potentials [1–29]. However, behaviors and internal states can be encoded across multiple neural modalities that measure different spatiotemporal scales [30–53], from small-scale spiking activity to field potentials which measure large-scale brain network activity [54–56]. For example, it has been shown that spiking and local field potential (LFP) activities exhibit shared dynamics, which are dominantly predictive of behavior during naturalistic reach-and-grasp movements [49]. Thus, building dynamical models that simultaneously incorporate multiple observation modalities is important for revealing how different spatiotemporal scales of neural population dynamics explain behavior. Further, such modeling can aggregate information across different spatiotemporal scales of neural activity to improve the performance of brain-machine interfaces (BMIs) [43, 44, 48–50]. We refer to dynamical modeling with multimodal observations as multiscale dynamical modeling.

Learning a multiscale dynamical model is challenging because different modalities have different statistical properties [43, 48, 49, 54–56]. For example, spike counts are discrete-valued action potential events with millisecond time-scales that are modeled well with Poisson distributions. In contrast, field potentials are continuous-valued and their spectral features evolve at slower time-scales than spikes, are extracted with slower time-steps, and are typically modeled with Gaussian distributions [41, 43, 44, 48, 49, 57–59]. To enable modeling of multimodal neural activity, we recently developed a multiscale expectation maximization (multiscale EM) method that learns a multiscale dynamical model by maximizing the joint log-likelihood of Poisson-Gaussian observations iteratively [48, 49]. However, EM is computationally expensive given its iterative numerical approach, which can be burdensome or even prohibitive especially for future real-time or adaptive learning applications (see Discussion). Thus, there is an important need for novel computationally efficient learning methods for multimodal neural data. Further, to enable real-time multiscale decoding applications, in addition to computational efficiency in learning, such novel methods should also produce models that support causal statistical inference from multimodal neural activity, which can be difficult to achieve (see Discussion).

Here, we develop an unsupervised learning method for multimodal Poisson-Gaussian data that is both computationally efficient and enables causal multiscale inference to fuse information across data modalities during decoding [60]. We also demonstrate the application of this method on multimodal spike-LFP neural activity recorded from the primate brain. To achieve computational efficiency, we develop a novel analytical method for learning multiscale dynamical models. This method extends the subspace identification (SID) techniques which currently only support single modalities to multimodal data [23, 26, 29, 61–65]. Importantly, our method also introduces a novel approach for ensuring the validity of the learned noise statistics, which is critical for enabling statistical and causal inference of the latent states from multimodal data after learning is completed. We term this method multiscale SID.

To date SID algorithms have been extended in various ways [23, 26, 29, 61–65], but not for addressing multiscale modeling. Traditional SID algorithms that model continuous signals operate by extracting the model parameters from empirically estimated cross-covariance matrices of future and past signals [19, 61, 62, 66]. SID has also been extended to two continuous signal sources to model the shared dynamics between continuous neural and behavioral signals [23], and for modeling the effect of input on neural-behavioral dynamics to dissociate input-driven and intrinsic neural dynamics [29]. Extensions of SID have also been developed for modeling discrete spike counts alone [63]. However, no SID method has been developed for joint dynamical modeling of multimodal observations that are a mix of continuous and discrete signals with different statistical properties.

To develop the multiscale SID method, we write a multiscale dynamical model with latent states and simultaneous discrete-continuous observations, e.g., consisting of spike counts and field potentials [48, 49]. We model the continuous observations as a linear Gaussian model of the latent states and the discrete spike counts as Poisson observations with a latent log firing rate that is a linear function of the same latent states. Extending the traditional SID to learn the parameters of this multiscale dynamical model involves several challenges.

The first challenge is related to the latent nature of the log firing rates [63, 67]. This latent nature means that the direct empirical estimation of the cross-covariance between the log firing rates and field potentials – which is needed by SID algorithms – is not possible. To address this challenge, we use statistical moment transformation [63, 68] and combine it with our multiscale dynamical model. Doing so, we find the multimodal cross-covariance between the latent log firing rates and the continuous modality indirectly by transforming the statistics that are directly computable from multimodal discrete-continuous observations.

The second challenge is to learn the model parameters while enforcing the learning of valid noise statistics. Learning valid noise statistics is not only important for accurate modeling, but also essential for statistical inference of latent states from neural observations, prediction of future neural activity, and neural decoding of behavior. Current covariance-based SID algorithms – i.e., those that can learn model parameters purely from data covariances as we do here – cannot guarantee learning of valid positive semidefinite (PSD) noise covariances [61–63]. Moreover, they cannot ensure that noise statistics conform to the multiscale dynamical model and its inference structure [43, 48]. The challenge of guaranteeing valid learned noise statistics remains unresolved even for the single-modal extension of SID for spike counts alone [63]. We address this challenge by devising a novel constrained optimization problem that revises the learned parameters of covariance-based SID methods to enforce valid noise statistics. We show that the model parameters learned by multiscale SID algorithm can then be used for causal multiscale inference of states, neural activity, and behavior.

Finally, multimodal observations may also be sampled at different rates, e.g., LFP spectral features often have a smaller sampling rate than binned spike counts [43, 48, 49], posing a challenge for jointly leaning and describing their dynamics given these different timescales. We show that in our datasets, this challenge can be addressed via an interpolation approach in the training data during model learning, enabling multiscale SID to jointly learn the dynamics of both modalities, even if they are sampled at different rates. After model is learned, inference can be done without interpolation.

We validate the multiscale SID algorithm in numerical simulations and on motor cortical spike-LFP recordings of a non-human primate (NHP) performing a naturalistic three dimensional (3D) reach-and-grasp movement task [23, 49, 69]. We find that multiscale SID can accurately learn the multiscale dynamical model parameters. In addition, we find that adding spiking signals to field potential signals or vice-versa improves the identification accuracy of dynamical modes and prediction of behavior compared to using single-modal activity, showing that multiscale SID can accurately fuse information across modalities. We also compare the multiscale SID to the recent multiscale EM algorithm [48, 49] in terms of accuracy and the time it takes to compute the model parameters, i.e., the computation time. In both simulations and the NHP dataset, we find that computation time required for the multiscale EM to converge was much higher than multiscale SID (about 180 times higher in simulations, and 30 times higher in the NHP dataset). Interestingly, this faster computation time for multiscale SID did not lead to degradation of accuracy. Indeed, for some metrics such as dynamical mode identification and neural prediction in our NHP dataset multiscale SID outperformed multiscale EM and in other metrics they performed similarly.

Taken together, multiscale SID provides an accurate and efficient method for learning multiscale dynamical models for multimodal neural population data while also enabling causal statistical inference from multimodal data. These new capabilities are especially important in real-time applications such as closed-loop or adaptive neuroscience studies or BMIs.

## 2. Results

### 2.1. Overview of multiscale SID

The derivation of multiscale SID is described in Methods and a detailed outline is provided in Algorithm 2. Here we provide a high-level overview of the multiscale SID algorithm (Algorithm 1, Figure 1b). We write a multiscale dynamical state-space model with latent state 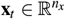, to jointly model the dynamics of continuous field potentials and discrete spike counts as

**Figure 1.**
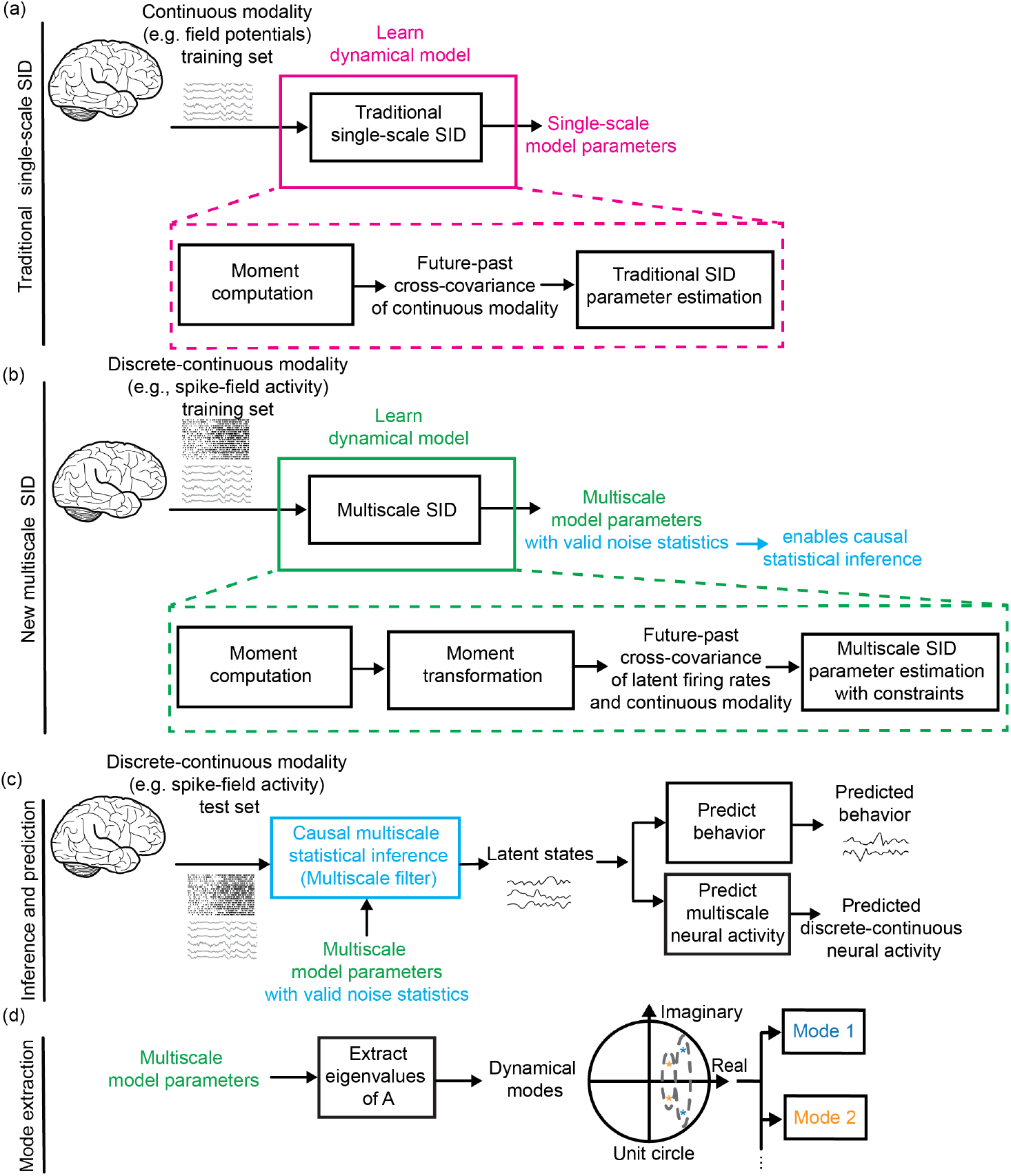
Multiscale SID algorithm for modeling and prediction of neural activity and behavior. **(a)** The traditional covariance-based SID algorithm learns the single-scale model parameters from a continuous modality (e.g., field potentials) in the training set (magenta box). These parameters are extracted from the future-past cross-covariance of the continuous modality **H**_*y*_, which can be computed directly from its observations. **(b)** The new multiscale SID algorithm (see Algorithm 1 and Algorithm 2) learns the multiscale model parameters from the discrete-continuous modalities (e.g., spike-field activity) in the training set (green box). Given that firing rates are latent, future-past cross-covariance of log firing rates and continuous modality **H**_*w*_ is not directly computable from the multimodal observations. Instead, we compute this cross-covariance **H**_*w*_ by transforming the moments of the discrete-continuous observations using the multiscale model equations. Then, we estimate the multiscale model parameters from **H**_*w*_ via SID methods. However, covariance-based SID methods even for a single modality do not guarantee valid noise statistics. We address this challenge by imposing added constraints in our SID method to enforce valid noise statistics within a novel optimization formulation. These constraints are critical for enabling multiscale statistical inference of latent states. **(c)** The learned valid parameter set is used to infer the latent states using a multiscale filter in the test set. These states are then used to predict behavior and the discrete-continuous neural activity using the learned model. **(d)** After learning the model parameters, the dynamical modes corresponding to a pair of complex conjugate eigenvalues or a real eigenvalue of **A** are computed.

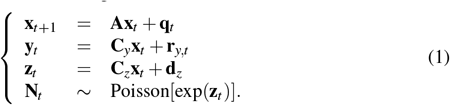

Here the continuous modality (e.g., field potential activity) denoted by 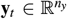 is modeled as a linear function of the latent states **x**_*t*_. The discrete spike counts denoted by 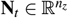 are modeled as Poisson-distributed with latent log firing rate 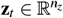, which is a linear function of the same latent states. The temporal dynamics of the latent state **x**_*t*_ are described using a linear state equation with a state transition matrix 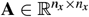. **q**_*t*_ and **r**_*y,t*_ are uncorrelated Gaussian noises that specify latent state and continuous modality measurement noises, respectively. 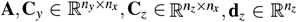, the state noise covariance 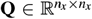 and the measurement noise covariance 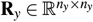 are the multiscale dynamical model parameters to be learned.

We recently developed an unsupervised multiscale EM algorithm [48, 49] to learn the multiscale model parameters in equation 1. However, this algorithm consists of iterative numerical steps and thus can require a long computation time for convergence. Here, we aim to develop a completely different analytical learning algorithm based on SID, which would enable computationally efficient learning of the model parameters in equation 1 while also enabling causal statistical inference of the latent states after learning. Such computationally efficient learning is not only desirable in general, but also essential for applications such as real-time adaptive BMI learning or for real-time tracking of plasticity in neuroscience experiments (see Discussion).

So far, SID algorithms or their extensions have not been able to model multimodal observations. Instead, SID methods have focused on modeling either only continuous modalities [19, 23, 26, 64, 65] or only discrete spike counts [63]. Here, we develop a multiscale SID algorithm that can learn a dynamical model for combined discrete-continuous observations. We also consider the case where these observations may also potentially be sampled at different rates. In what follows, we briefly mention the main challenges involved and how we solve them to develop the multiscale SID algorithm. Details are in the Methods section.

The first challenge is that unlike continuous signals (e.g., field potentials), the log firing rates of spiking activity are not directly measurable. Traditional single-scale SID algorithms [61, 62] rely on measured continuous signals to compute the cross-covariance between the future and past of these signals 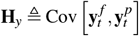 (Figure 1(a)). The cross-covariance is then used to estimate the single-scale model parameters in the following equation:

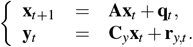

where future and past vectors 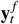 and 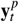 are formed by stacking time-lagged continuous signals (equations 8 and 9 in Methods). Extension of SID to single-modal Poisson spike counts has computed this covariance for the latent log-firing rates by transforming the moment of the associated observable spikes [63]. In comparison, in our multiscale model (equation set 1), we have multimodel discrete-continuous observations. Thus, we need to compute the cross-covariance of future and past of a signal that is the concatenation of the latent log firing rates as well as the continuous observation modality, **H**_*w*_ (equation 14). However, since log firing rates **z**_*t*_ are latent, direct empirical computation of the cross-covariance matrix elements that involve the log firing rates **z**_*t*_ is not possible. To address this challenge, we compute these elements via a transformation of the joint spike count and continuous signal statistics (i.e., moments), which are directly computable from observed samples of multimodal discrete-continuous neural data **N**_*t*_ and **y**_*t*_ (see steps 1-2 of the Algorithm 2, section 4.2.3, and [63]). Note that given the latent nature of log firing rates, we had to use a covariance-based SID approach (see Discussion).

The second challenge is that covariance-based SID algorithms cannot guarantee that the learned noise statistics will have valid PSD covariances and will conform to the multiscale dynamical model and inference structures in equation 1 [43, 48], which have the following requirements: 1) state and continuous observation noise covariances, **Q** and **R**_*y*_, are valid covariances (i.e., are PSD), 2) the log firing rates **z**_*t*_ do not have an additive noise term, and 3) state and continuous observation noises, **q**_*t*_ and **r**_*y,t*_, are uncorrelated. Valid noise statistics are important both for accurate modeling and for enabling statistical inference of latent states after learning. This challenge of identifying valid noise statistics is also faced by the extension of covariance-based SID methods for single-modal spike counts [63], but remains largely unaddressed even in that case. To address this challenge, we formulated an optimization problem that enforces the above three requirements (equation 39, section 4.2.5). We then solved this optimization with semidefinite programming (see step 9 of the Algorithm 2 and section 4.2.5, [70–72]).

Having addressed the aforementioned challenges, we developed the multiscale SID algorithm that identifies the multiscale model parameter set *𝒩* = {**A, C**_*y*_, **C**_*z*_, **Q, R**_*y*_, **d**_*z*_} from multimodal neural observations **N**_*t*_ and **y**_*t*_ (Figure 1(b)). We provide a summary of the main steps of multiscale SID here in Algorithm 1 (for more details see Algorithm 2 and section 4.2.5).

#### Algorithm 1 Multiscale SID summary

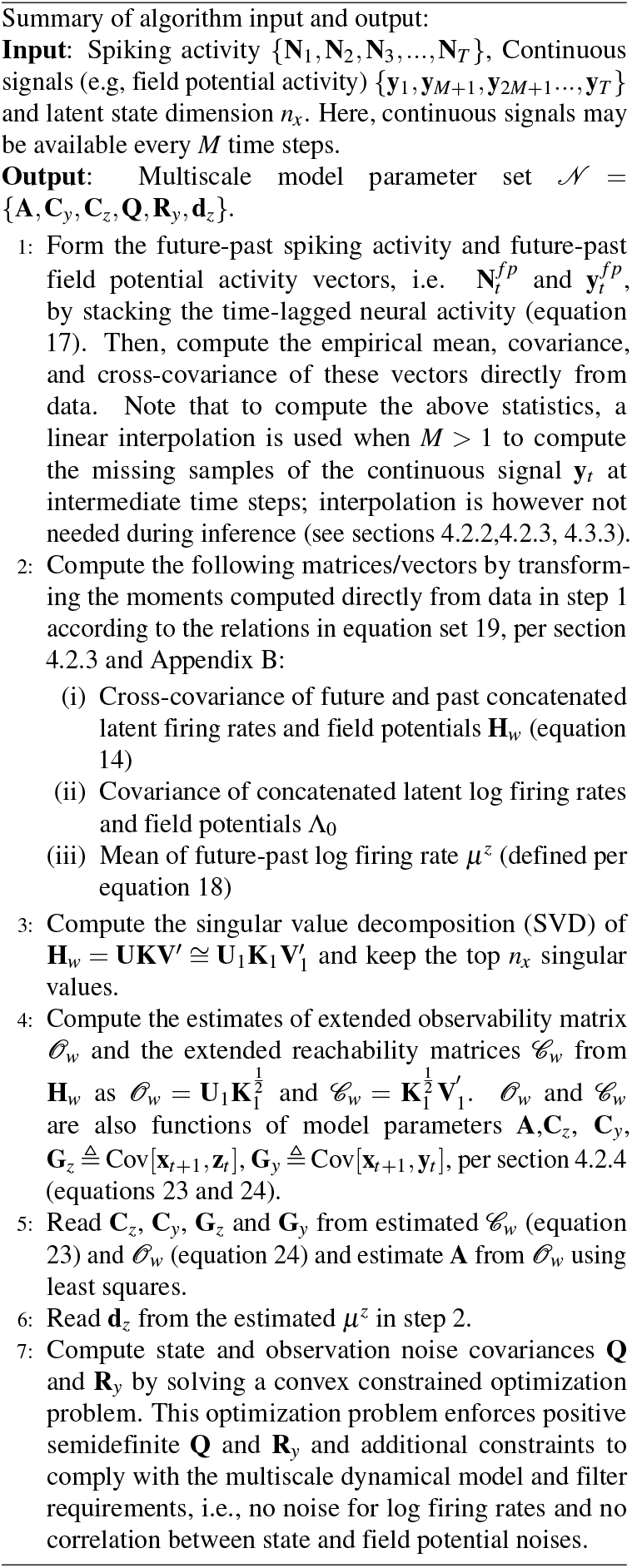

Because the multiscale SID algorithm ensures valid noise statistics and conforms to the requirements of the multiscale dynamical model/inference structures, its learned parameters can readily enable statistical and causal inference of latent states from multimodal neural data. This can be done by incorporating the learned parameters inside a multiscale filter [43]. The inferred latent states can also be used to predict neural activity and behavior.

### 2.2. Simulation validations: Multiscale SID performs accurately while being substantially more computationally efficient

We simulated multimodal discrete-continuous neural data from models with random parameters according to equation 1 (see section 4.3.1 for simulation details). We then applied the learning algorithms to the simulated data to learn the model parameters, identify dynamical modes, extract the latent states and predict neural activity (see sections 4.2.5, 4.3.3, Figure 1(b)-(d)). Each dynamical mode corresponds to a pair of complex conjugate eigenvalues or a real eigenvalue of the state transition matrix **A**. To show that multiscale SID in Figure 1(b) can successfully aggregate multimodal data, we compared it with single-scale SID algorithms for continuous field potentials alone [61] and for discrete spikes alone [63]. To show that multiscale SID achieves good accuracy while being substantially more computationally efficient, we compared it with the existing multiscale EM algorithm for multimodal spike-field neural data [48, 49]. Performance measures are detailed in section 4.3.2-4.3.3. Note that while we refer to the discrete-continuous data as spike-field for ease of exposition, these are general simulated multimodal data; thus, these simulations validate multiscale SID broadly for multimodal Poisson and Gaussian data observations.

#### 2.2.1. Multiscale SID accurately identifies the model parameters

We found that model parameters were identified with decreasing normalized error as the number of training samples increased (Figure 2). All model parameters could be identified accurately, with the normalized error reach-ing below 6% when trained with *T* = 10^6^ training samples (Figure 2).

**Figure 2.**
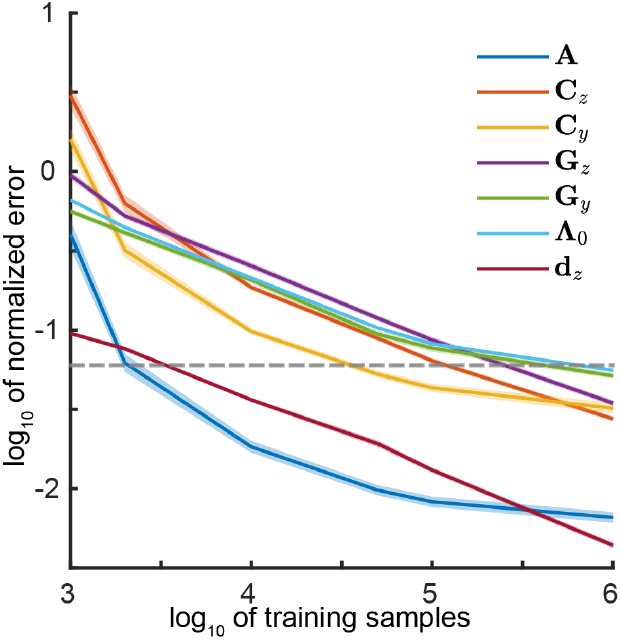
Multiscale SID accurately identifies the model parameters in simulations. Normalized identification error of all parameters in multiscale SID as a function of number of training samples across 50 randomly generated multiscale models. All parameter identification errors decrease as more training samples are used. Using 10^6^ samples, all normalized errors are less than 6%. The dashed horizontal line indicates 6% normalized error. Solid lines show the mean and shaded area represents s.e.m. The set {**A, C**_*z*_, **C**_*y*_, **G**_*z*_, **G**_*y*_, Λ_0_, **d**_*z*_} fully characterize the multiscale model in equations 2, where 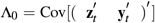 is the covariance of concatenated log firing rates and field potential signals, and **G**_*y*_ = Cov[**x**_*t*+1_, **y**_*t*_] and **G**_*z*_ = Cov[**x**_*t*+1_, **z**_*t*_] are the cross-covariances of latent states with log firing rates and field potential signals, respectively (see sections 4.2.5, 4.3.2).

#### 2.2.2. Multiscale SID outperforms multiscale EM in computation time and in identification of dynamical modes, while reaching a similar neural prediction accuracy

For the same 50 simulated systems as in section 2.2.1, we compared the computation time of multiscale SID with that of multiscale EM in learning the model parameters as well as their performance in identifying dynamical modes and prediction of neural activity. We continued the multiscale EM iterations until convergence in dynamical mode identification error or up to 175 iterations, whichever happened earlier (see sections 2.2, 4.3.4). We refer to this learnt model as converged EM and use it for comparison of EM and SID performance. We compute and compare the performances as a function of training sample size (Figures 3 and 4). We also separately highlight the results for *T* = 5 × 10^4^ training samples, which is in the same order of magnitude as the session lengths of the NHP dataset used in this study (2.79 × 10^4^ ± 0.14 × 10^4^ (mean ± s.e.m.) samples).

**Figure 3.**
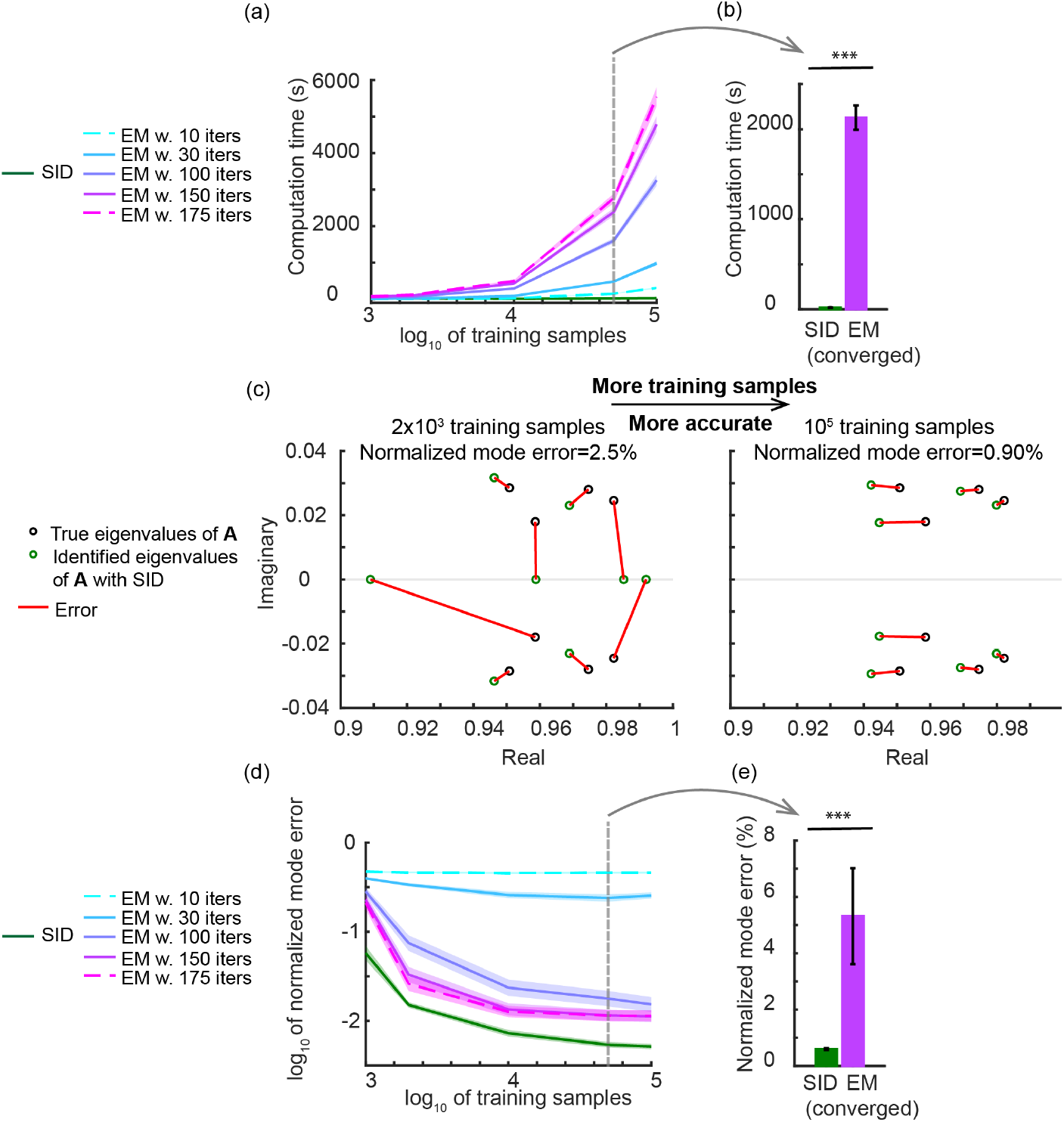
Multiscale SID outperforms converged multiscale EM in computation time and identification of dynamical modes in simulations. (**a**) Computation time (in seconds) of multiscale SID and multiscale EM with different number of iterations as a function of number of training samples. Multiscale SID has a much lower computation time compared with multiscale EM. Also, the computation time of multiscale EM monotonically increases with more iterations or training samples. Solid line represents the mean across 50 simulated models and shaded area represents s.e.m. (**b**) Computation time of multiscale SID vs converged multiscale EM using 5 × 10^4^ training samples (indicated by vertical dashed line on panel (a)). This training sample size has the same order of magnitude as the NHP dataset. The bar represents the mean and the whiskers indicate s.e.m. Asterisks indicate significance of performance comparison between multiscale SID and converged multiscale EM, i.e., multiscale EM for the iteration at which mode error converges. (*: P *<* 0.05,**: P *<* 0.005,***: P *<* 0.0005) (**c**) For one simulated multiscale model, the true and SID identified eigenvalues of the state transition matrix **A** are shown in black and green circles for 2 × 10^3^ (left) and 5 × 10^4^ training samples (right). Red lines indicate the identified eigenvalue errors, that is mode errors. Each dynamical mode corresponds to a pair of complex conjugate or a real eigenvalue of the true state transition matrix **A** (Figure 1(d)). Normalized mode error is computed by first dividing the sum of the squared length of the red error lines by the sum of the true eigenvalue squared magnitude, and then taking its square root (section 4.3.2). Normalized mode error decreases with increasing the training sample size. (**d**)-(**e**) Same as (b)-(c) but for normalized mode error. Multiscale SID has a significantly lower mode error compared with multiscale EM. The normalized mode error monotonically decreases with more iterations for multiscale EM and with more training samples for both multiscale EM and SID.

**Figure 4.**
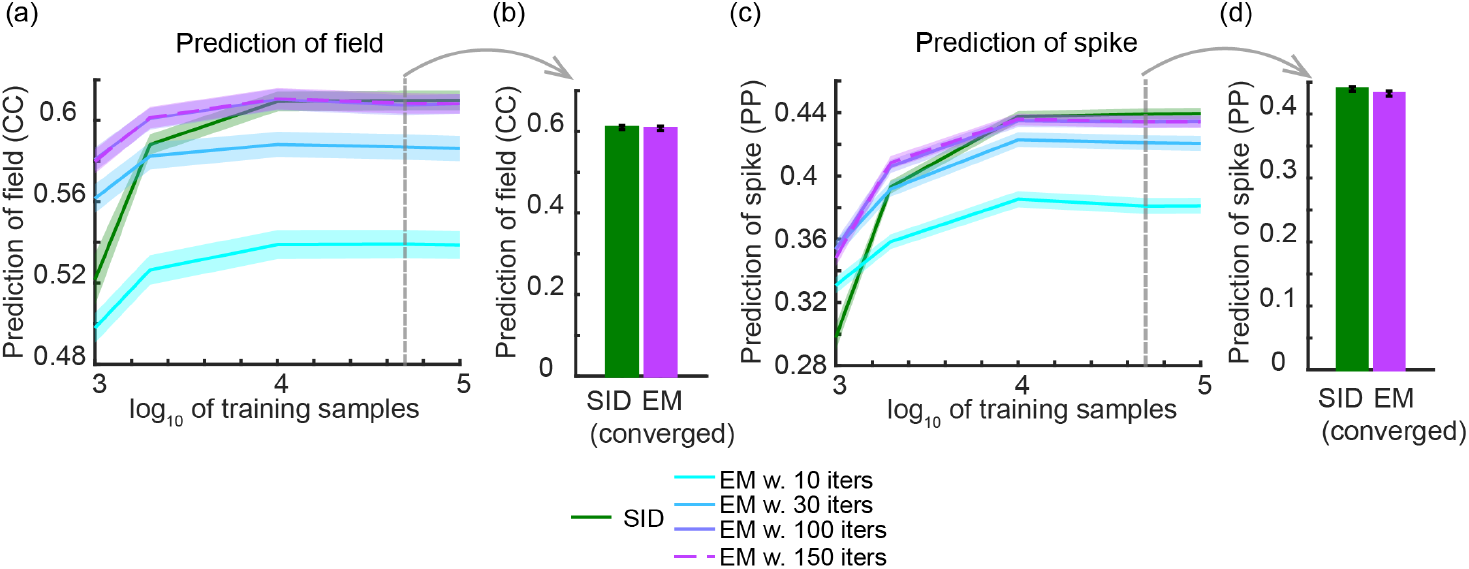
While being significantly more computationally efficient and having better dynamical mode identification (Figure 3), multiscale SID is also as accurate as converged multiscale EM in predicting spiking and field potential data in simulations. (**a**) One-step-ahead prediction accuracy of field potential activity quantified by correlation coefficient (CC) between the one-step-ahead predicted and the true field potential activity for multiscale SID and multiscale EM (with different number of iterations) as a function of training samples. (**b**) One-step-ahead prediction performance of field potential activity for multiscale SID vs converged multiscale EM using 5 × 10^4^ training samples, which is similar to the number of samples in the NHP datasets and indicated by the vertical dashed line on panel (a). (**c**)-(**d**) Same as (a)-(b) but for one-step-ahead prediction accuracy of spiking activity quantified by prediction power (PP). Figure convention is as in Figure 3. Overall, multiscale SID performs similarly in neural prediction to multiscale EM in simulations when enough training samples—but still comparable to that of the real data—are available.

##### Computation time

First, the computation time required for learning in multiscale SID was orders of magnitude faster than converged multiscale EM (Figure 3(a)-(b)). For example, for *T* = 5 × 10^4^ training samples, the learning time for multiscale SID was 18.85 ± 2.16 s vs. 2127.80 ± 133.70 s for converged multiscale EM, i.e., 186.00 ± 20.41 times faster (Figure 3(b), *P* = 3.89 × 10^−10^, *N*_*s*_ = 50). Further, the computation time of multiscale EM grew roughly exponentially with training sample size, while the computation time of multiscale SID had minimal changes as the training sample size increased (Figure 3(a)). Also, as expected, more EM iterations required increasingly more time to run (Figure 3(a)). These results indicate that multiscale EM is increasingly more computationally expensive to run compared with multiscale SID for larger training sample sizes. Overall, multiscale SID was significantly faster than the converged multiscale EM across all simulated training sample sizes (FDR-corrected *P*≤ 2.40 × 10^−4^, *N*_*s*_ = 50).

##### Dynamical modes

Second, we explored the accuracy in identifying the dynamical modes, which, as noted earlier, are the eigenvalues of the state transition matrix **A** and quantify the dynamics. For visualization, we show one example simulated model in Figure 3(c) that illustrates the error between the true and identified eigenvalues from which the normalized mode error is computed per section 4.3.2 and how this error is decreased by increasing training sample size. Interestingly, multiscale SID was also significantly more accurate than the converged multiscale EM in identifying the dynamical modes (Figure 3(d)-(e)). For *T* = 5 × 10^4^ training samples, the normalized mode error in multiscale SID was 0.60 ± 0.04% vs. 5.31 ± 1.70% for converged multiscale EM (Figure 3(e), *P* = 3.99 × 10^−8^, *N*_*s*_ = 50).

Even as we increased the training sample size, mode identification in multiscale SID was significantly more accurate than the converged multiscale EM across all simulated training sample sizes (FDR-corrected *P*≤ 3.20 × 10^−3^, *N*_*s*_ = 50). Also, as expected, the normalized mode error monotonically decreased with increasing training sample size for both algorithms, and with more EM iterations for multiscale EM (Figure 3(d)). The reasons for more accurate performance of multiscale SID in mode identification could be the approximations that must be made in multiscale EM to find the posterior density, the fact that EM aims to optimize the neural data likelihood rather than dynamic mode identification, and that multiscale EM does not guarantee to even optimize the neural data likelihood given its approximations [6, 73] (see Discussion).

##### Neural prediction

Third, we compared the multiscale SID and multiscale EM in predicting the simulated multimodal neural activity (see sections 4.3.3, Figure 4). We found that despite being much faster (Figure 3(b)) and while enabling more accurate dynamical mode identification (Figure 3(e)), multiscale SID had comparable performance to the converged multiscale EM even in neural prediction when provided with enough training samples (Figure 4). For field potentials, this accuracy was quantified with correlation coefficient (CC) between the predicted and true field potential activity and for spiking activity with prediction power (PP) (defined in section 4.3.3). With *T* = 5 × 10^4^ training samples, the one-step-ahead prediction accuracy of field potentials and spiking activity for multiscale SID and converged multiscale EM were within 0.38% and 1.7% of each other, respectively (Figure 4(b),(d)). Also, the prediction of neural activity monotonically improved with training sample size for both algorithms and with EM iterations for multiscale EM (Figure 4(a),(c)).

#### 2.2.3. Multiscale SID can fuse information across discrete and continuous neural modalities and identifies the dynamical modes better than single-scale SID

Multiscale modeling allows information across multiple neural modalities to be aggregated and thus has the potential to outperform modeling of any single modality in terms of learning the neural dynamics. To demonstrate this capability, we simulated 50 multiscale models with 14 discrete spiking signals, 14 continuous field potential signals and 4 dynamical modes. We then gradually added increasingly more neural signals from one modality to a fixed number of signals from the other modality, referred to as the baseline signals (section 4.3.5). We used single-scale SID to identify a model for the baseline signals and used multiscale SID to do so for the combination of baseline signals with the signals from the second added modality.

We found that the learned models became increasingly more accurate as more and more signals of a second modality were added to the signals from the first baseline modality (Figure 5(a),(c)). Specifically, the normalized mode error monotonically decreased as field potential signals were added gradually to a fixed number of spiking signals or vice versa (Figure 5(a),(c)). These results suggest that multiscale SID can correctly aggregate information across discrete and continuous modalities.

**Figure 5.**
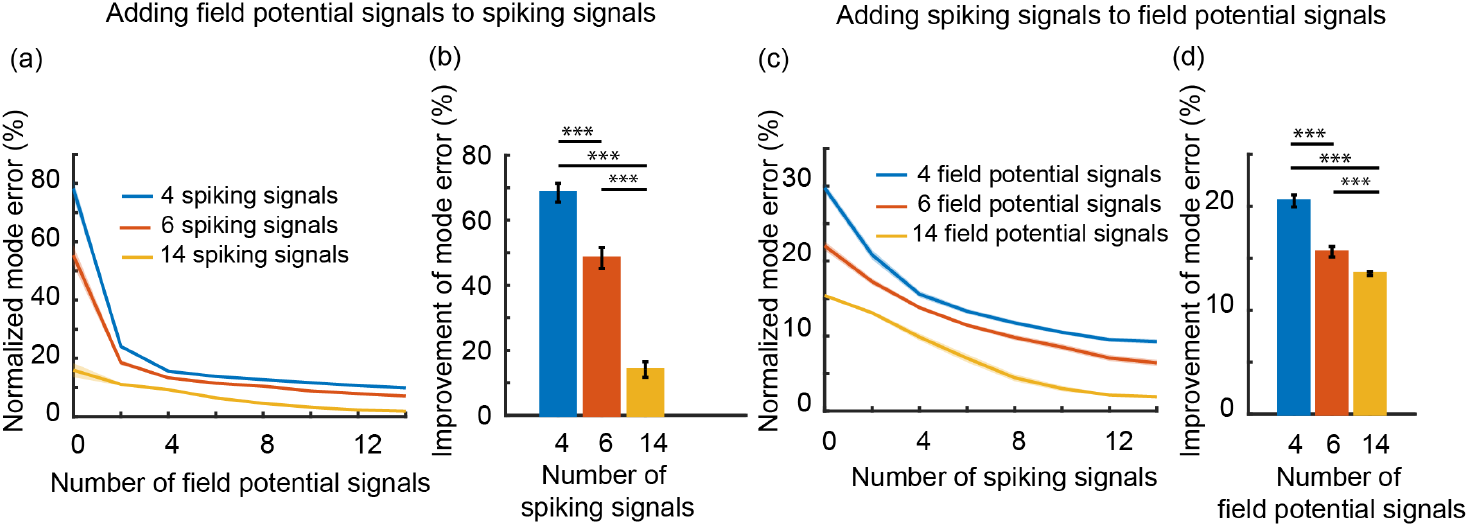
Multiscale SID outperforms single-scale SID in identification of dynamical modes in simulations due to fusion of information across modalities, with the largest performance gain being obtained in the low information regime. **(a)** Normalized mode error as continuous field potential signals are added gradually to 4, 6, or 14 discrete spiking signals. The start of the curves (i.e., 0 on x-axis) indicates normalized mode error for single-modal signals (i.e., spiking signals only). Solid line indicates mean across 50 simulated neural network systems and shaded area represents s.e.m (*N*_*s*_ = 50 data points). **(b)** Comparison of the improvement of normalized mode error after adding 14 continuous field potential signals to 4, 6, or 14 discrete spiking signals. Bars indicate mean and whiskers show s.e.m. Asterisks indicate significance of a pairwise comparison of improvement value across baseline regimes (*: P *<* 0.05,**: P *<* 0.005,***: P *<* 0.0005). **(c)-(d)** Same as (a)-(b) but for adding spiking signals to field potential signals.

We next compared the improvement gained by going from single-scale to multiscale modeling for cases with different numbers of baseline signals (Figure 5(b),(d)). We found that the improvement in identification error of dynamical modes was larger for cases with fewer number of baseline signals (Figure 5(b),(d), FDR-corrected *P* ≤ 1.64 × 10^−4^, *N*_*s*_ = 50). This result suggests that for learning the dynamics, multiscale modeling has the most benefit compared with single-scale modeling in the low information regime, i.e., when the initial modality has incomplete information about the neural dynamics.

In these numerical simulations and motivated by previous studies [49], we simultaneously simulated modes that were shared between the two modalities as well as modes that were exclusive to each of the modalities (distinct modes) (see section 4.3.1). We found that the addition of one modality to another improved the learning of both the modes that were exclusive to the added modality as well as the modes that were shared between the two modalities. This result was found by analyzing the identification error of distinct and shared modes separately (Figure A1), and again shows that information is being aggregated across modalities about their collective dynamics to learn them more accurately.

### 2.3. Multiscale SID accurately predicts the spike-LFP recordings from the NHP brain during naturalistic movements, while being substantially faster

We next used multiscale SID to model multimodal spiking and LFP activity recorded from an NHP while performing a naturalistic 3D reach and grasp movement task (Figure 6(a), see section 4.4.1 and [23, 49, 69] for more details). We obtained discrete spiking activity **N**_*t*_ by detecting the threshold crossing events every 10 ms and field potential activity **y**_*t*_ by computing power features in seven standard frequency bands from the recorded neural signals every 50 ms (see section 4.4.2). During model learning alone, we interpolate the power features to recover the samples of **y**_*t*_ at every 10 ms time-step that spike counts are observed; note, during inference, no interpolation is necessary and these intermediate field samples can simply be treated as missing observations in a multiscale filter [43] (see sections 4.2.2,4.2.3,4.3.3). We used a five-fold cross-validation scheme. We learned the multiscale model using multiscale SID in the training data and then used it in the test data to extract the latent states and predict the spike-LFP activity and behavior (i.e., joint angles) from the extracted latent states (see sections 4.3.3,4 We also compared the multiscale SID with the existing multiscale EM for spike-LFP data and with single-scale SID for spikes alone or LFP alone (sections 4.4.5,4 For each method, we fitted models with latent state dimensions spanning *n*_*x*_ ∈ {2, 4, …, 24} (section 4.4.5). Convergence criteria for EM was set based on convergence of neural prediction performance as described in section 4.4.5, with maximum allowed EM iterations of 150.

**Figure 6.**
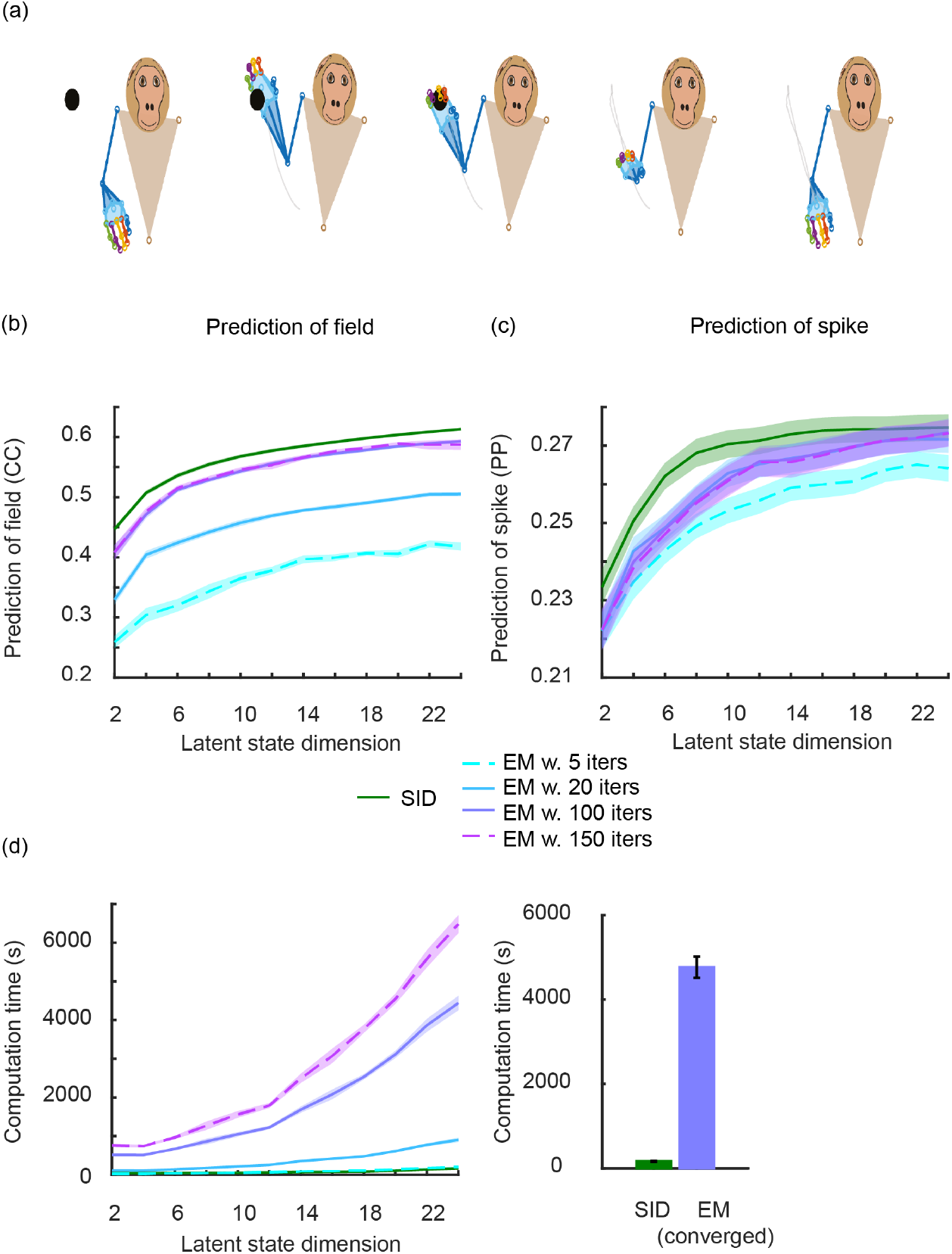
In NHP datasets, multiscale SID outperforms converged multiscale EM in both computation time and neural prediction accuracy. (**a**) Spike-LFP activity was recorded as a non-human primate performed naturalistic reach and grasp movements to randomly positioned objects in 3D space. (**b**)-(**c**) One-step-ahead prediction accuracy for field potentials and spiking activity as a function of latent state dimension (measured by CC and PP, respectively). Solid line represents the mean and shaded area represents s.e.m. over folds and data sessions (35 data points). (**d**) **Left**: Same as (b)-(c) but for computation time (in seconds). Computation time monotonically increases with more multiscale EM iterations or latent state dimensions. **Right**: Comparison of multiscale SID and converged multiscale EM at *n*_*x*_ = 24. Multiscale SID is 30 times faster than multiscale EM.

#### 2.3.1. Multiscale SID outperforms multiscale EM in computation time and spike-LFP prediction in the NHP dataset

We found that multiscale SID was significantly faster than the converged multiscale EM across all latent state dimensions (Figure 6(d) left panel, FDR-corrected P-value ≤ 1.29 × 10^−7^, N_*s*_ = 35). Interestingly, in addition to being faster, multiscale SID was also significantly more accurate in prediction of LFP across all latent state dimensions and in prediction of spiking activity across latent state dimensions up to 22 (Figure 6(b),(c), FDR-corrected P-value ≤ 1.70 × 10^−2^, N_*s*_ = 35). For example, for the maximum considered state dimension of *n*_*x*_ = 24, multiscale SID was 30.66 ± 1.84 times faster than the converged multiscale EM while reaching 3.5% more accurate LFP prediction CC and slightly more accurate spike prediction PPs (Figure 6). Thus, in addition to its substantially lower computational cost, multiscale SID could outperform or do comparably to multiscale EM. This result is consistent with our simulations in section 2.2.2 and again likely due to the approximations (e.g., Laplace) in multiscale EM that lead to inaccuracy (see Discussion).

Finally, we computed the accuracy in predicting behavior as quantified by the correlation coefficient (CC) between the predicted and true joint angle trajectories (see section 4.4.3). We found that despite its much faster training time, multiscale SID had similar accuracy in predicting behavior compared to the converged multiscale EM (Figure A2), and that it took multiscale EM substantially longer training time to achieve this accuracy. The substantially longer training time in multiscale EM is due to its iterative and computationally expensive nature, and meant that multiscale SID was able to reach better or comparable modeling accuracy using much faster computations (Figure 6(d)).

#### 2.3.2. Multiscale SID improved behavior decoding in the NHP dataset compared with single-scale SID due to addition of spike-LFP information

We next performed a neural signal addition analysis similar to what we performed for simulated data (sections 4.4.6, 4.3.5, 2.2.3). We constructed a pool of 30 spike channels and 30 LFP channels and gradually added signals from one modality to a fixed number of signals from the other modality, referred to as the baseline neural signals (section 4.4.6). We identified a model for the baseline neural signals on their own using the single-scale SID and learned models for the multimodal signals (baseline plus added signals) using the multiscale SID. We computed the cross-validated behavior prediction accuracy for each learned model.

We found that behavior prediction performance benefited from multiscale modeling (Figure 7), and monotonically improved both as field potential signals were added to baseline spiking signals and vice versa (Figure 7(a),(c)). Further, similar to our simulation results, the improvement in behavior prediction performance using multiscale SID was larger for the lower information regime, i.e., when the number of baseline signals was smaller (Figure 7(b),(d), FDR-corrected *P* ≤ 1.45 × 10^−27^, *N*_*s*_ = 350). Also, this improvement was obtained regardless of whether the baseline modality was the discrete spiking or the continuous LFP modality. This bidirectional improvement suggests that the advantage of multiscale over single-scale modeling was not simply due to the dominance of one modality over the other, but rather due to the addition of information across them. Together, these results suggest that for NHP multimodal spike-LFP data, multiscale SID is correctly aggregating information across neural modalities.

**Figure 7.**
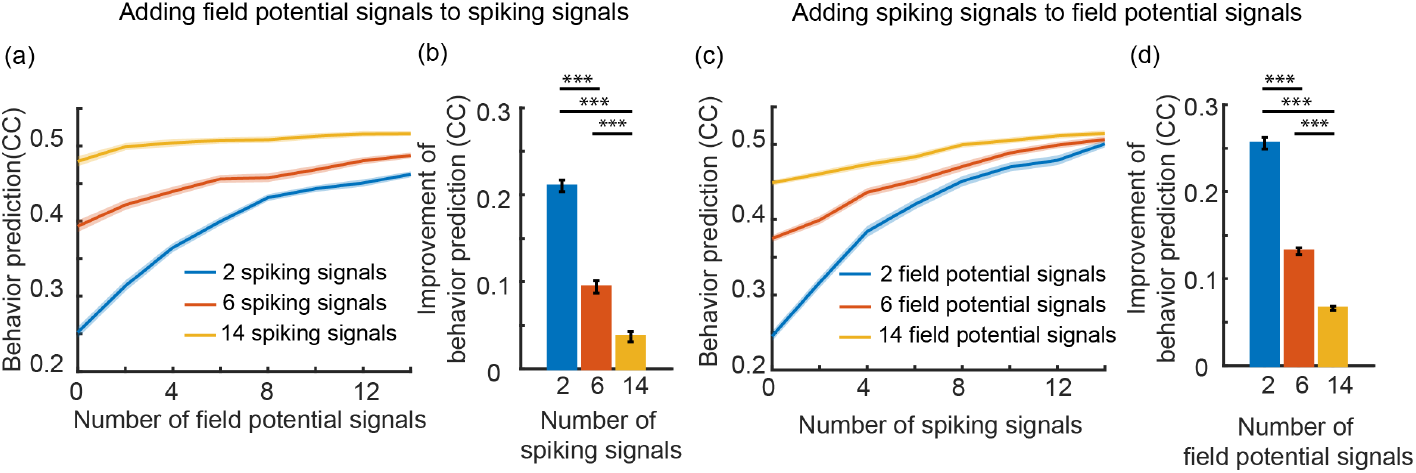
In NHP datasets, multiscale SID improves the behavior prediction accuracy compared to single-scale SID due to fusion of information across modalities and this improvement is largest in the low information regime. (**a**) Behavior prediction accuracy quantified by correlation coefficient (CC) between the predicted and true behavior as field potential signals are added gradually to 2 (blue), 6 (red) and 14 (yellow) spiking signals. The start of the curves (i.e. 0 on x-axis) indicates behavior prediction for single-modal neural activity (i.e. spiking signals only). Solid line indicates mean across folds, random sets of selected neural signals and data sessions and shaded area represents s.e.m (*N*_*s*_ = 350 data points). (**b**) Comparison of the improvement of behavior prediction (quantified by difference of CCs) after adding 14 field potential signals to 2 (blue), 6 (red) and 14 (yellow) spiking signals. Bars indicate mean and whiskers show s.e.m. Asterisks indicate significance of a pairwise comparison of improvement value (*: P *<* 0.05,**: P *<* 0.005,***: P *<* 0.0005). (**c**)-(**d**) Same as (a)-(b) but for adding spiking signals to field potential signals.

## 3. Discussion

We developed multiscale SID, an analytical method that efficiently learns multiscale dynamical models of multimodal discrete-continuous spike-field population activity, extracts their low-dimensional latent dynamics, and enables causal and multimodal statistical inference of latent states, as well as prediction of neural activity and behavior. We validated this method using extensive numerical simulations and NHP spike-LFP activity recorded during 3D naturalistic reach and grasp movements [49, 69]. Multiscale SID accurately learned the multiscale model parameters and the low dimensional dynamical modes in spike-field population activity. Also, it fused information across spiking and field potential modalities, thus more accurately identifying the dynamical modes and predicting the behavior compared to when using either modality alone. Moreover, multiscale SID had a much lower computational cost compared to the existing multiscale EM method, while being better in identifying the dynamical modes and having a better or similar accuracy in predicting neural activity and behavior. These capabilities are important in studying multimodal neural dynamics and developing multimodal neurotechnology especially for real-time and adaptive experiments. Finally, while we focused on modeling multimodal spike-field dynamics, multiscale SID provides a general analytical method that can be applied broadly to other multimodal discrete-continuous time-series.

### 3.1. A computationally efficient method to learn multiscale dynamical models and enable causal statistical inference

Developing methods that can identify a low-dimensional subspace for multiscale dynamics and model these dynamics is challenging because of the different statistical properties of multimodal signals. For example, spike counts are discrete valued with millisecond timescales while field potentials are continuously varying and their typically-studied spectral features evolve at a slower timescale and are usually sampled with at a slower rate than spikes [41, 43, 44, 48, 49, 54–59, 74].

We recently resolved this challenge by developing a multiscale Expectation Maximization algorithm (multiscale EM) [48] and successfully demonstrating it for modeling of spike-field neural data [48, 49]. However, EM has a high computational cost given its iterative approach, which can be burdensome or even prohibitive especially for realtime or adaptive learning applications. Here we addressed this challenge by developing an analytical computationally efficient multiscale subspace identification algorithm [60]. We showed that multiscale SID not only substantially reduced computation time, but also performed with similar or better accuracy compared with the existing multiscale EM method for multimodal spike-field data.

Beyond EM, other numerical techniques can also be computationally expensive and in many cases may not enable causal statistical inference. For example, prior works have developed non-causal numerical variational inference methods for continuous fMRI and categorical behavioral data [75]. But these techniques did not focus on spike-field neural data. Moreover, similar to EM, these methods can have a high computational cost compared with SID given their iterative numerical approach, which is burdensome especially for real-time or adaptive learning. Finally and critically, enabling real-time applications requires the ability for causal inference/decoding, which is not achieved by these variational inference methods [75]. The new multiscale SID method addresses all these challenges.

### 3.2. Challenges in developing a multiscale SID method

Prior subspace identification (SID) methods have been designed for modeling the dynamics in just a single modality of data. We had to address multiple challenges in developing the multiscale SID for multimodal data.

First, while spike counts (**N**_*t*_) are observable, their log firing rates (**z**_*t*_) which relate them to the latent states (**x**_*t*_) are latent (equation 2). For example, prior work for single-modal spiking activity have computed the covariance of log firing rates by transforming that of the observable spikes [63]. Since one of the modalities in our multiscale dynamical model is the spiking modality, we cannot directly estimate the cross-covariance of its log firing rates with the continuous modality (**y**_*t*_, e.g., field potentials). These cross-covariance terms between the two modalities are required for parameter learning. We address this challenge by deriving a mathematical transformation that computes these cross-covariances between the latent log firing rates and the continuous observations based on the covariances of the observed spike counts and continuous modality (section 4.2.3, equation set 19).

Second, another key challenge in developing multiscale SID was that noise statistics in the multiscale SID model (equation 1) must satisfy certain conditions including being positive semi-definite (PSD). These conditions are important not only for accurate modeling, but also, critically, for enabling statistical inference of the latent states and behavior. While there are traditional non-covariance-based SID methods that can at least guarantee PSD conditions for covariance matrices [61, 62], these algorithms are not applicable for the multiscale model (equations 1 and 3). This is because these methods require direct access to continuous observations—and not just their covariances—, but log firing rates (**z**_*t*_) are not observable and their corresponding observable spike counts are not continuous (see section 4.2.2). Thus, after computing the cross-covariances, we had to utilize a covariance-based SID approach, which does not guarantee valid noise statistics as is known in the literature [61]. To address this challenge, we introduced a novel approach where we devised a constrained optimization problem to learn valid noise statistics (section 4.2.6). Beyond the positive semi-definiteness of noise covariances, this flexible constrained optimization approach further allowed us to incorporate other conditions needed for conforming to the multiscale model and its inference structure (e.g., **R**_*z*_ = 0 in equation 3). Flexibly imposing such additional constraints is not addressed in current covariance-based or non-covariance-based SID methods [61–63]. Our novel constrained optimization approach could be used by future work in other settings involving SID methods and their extensions, for example in developing SID for other observation distributions or to impose alternative constraints on noise statistics learned by SID.

Finally, different modalities may be sampled at different rates. For example, spike counts typically have a faster sampling rate than LFP spectral features [43, 48, 49], which was also the case in our NHP data (see Methods for details). We showed that in our datasets, this challenge can be addressed with an interpolation approach in the training set, enabling multiscale SID to learn the collective dynamics of the modalities even if they are sampled at different rates. Once model parameters are learned, we can process the multimodal data with different sampling rates without interpolation during multiscale inference [43].

### 3.3. Comparison of multiscale SID with EM in computation time and accuracy

While multiscale EM learns a set of parameters for the same model structure, our results in simulations and in real data show that multiscale SID has much lower computation cost while achieving better dynamical mode identification and better or similar neural prediction. Moreover, this advantage gets more pronounced as the training sample size increases as shown in Figure 3(a),(d). There could be multiple reasons for this.

In terms of computation cost, the higher computation time of multiscale EM is because of its iterative numerical nature. At each iteration, multiscale EM needs to run the expectation step which involves filtering and smoothing the entire training data [43, 48] and the maximization step, which requires solving an optimization problem [48]. In contrast, multiscale SID largely consists of a specific set of non-iterative analytical algebraic operations.

In terms of accuracy, our results show that multiscale SID learns the dynamical modes more accurately than multiscale EM, and performs better or similarly for neural prediction in real neural data or in simulations when provided with enough training samples (Figures 3(d)-(e), 6(b)-(c)). The better performance of multiscale SID could be due to the approximations that multiscale EM has to make to find the posterior density. In particular, multiscale EM uses the Laplace approximation to approximate the posterior density with a Gaussian distribution, just as is done in single-scale EM for Poisson observations [4, 76]. Given that the Laplace approximation may fail to capture broader statistics of the true posterior distribution [77], it may lead to suboptimal parameter identification and neural predictions. Furthermore, given the Laplace approximation, there is no guarantee for non-decreasing data likelihoods in consecutive iterations, which is the objective of EM [6, 73].

Overall, the higher efficiency of multiscale SID while maintaining better or similar accuracy can make it beneficial for multiscale modeling especially when efficient learning is desired or needed.

### 3.4. Multiscale SID for other multimodal distributions

Here we derived the multiscale SID for joint modeling of continuous Gaussian and discrete Poisson modalities (equation 1). However, the multiscale SID framework can flexibly generalize to other multimodal distributions as long as they conform to the generalized linear models (GLMs). This can be done because the moment transformation step for empirical estimation of covariances **H**_*w*_ is flexible and easily extendable to other GLMs, and because the other steps of the derivation do not depend on the distribution of observations, and the constrained optimization here can flexibly enforce assumptions needed for different observation distributions. Also, while our demonstrations in real data were for spike-LFP recordings, future work can apply the multiscale SID for modeling of spikes along with other continuous neural modalities such as intracranial EEG and electrocorticogram (ECoG). More broadly, beyond neural data, multiscale SID can also be applied to other multimodal discrete-continuous time-series to model them collectively.

### 3.5. Information fusion across neural modalities

Different neural modalities measure distinct spatiotemporal scales of information in the brain, and thus may contain both shared and distinct information about various brain states. Allowing for multimodal modeling can not only improve the decoding of shared information, but also allow for decoding of distinct information that may not be possible with a single modality. This capability is enabled because multiscale SID learns not only the dynamics that are shared across the modalities, but also dynamics that are distinct to either modality, i.e., the collective dynamics of both modalities. Consequently, multimodal modeling can also make future neurotechnologies more robust to neural signal loss. For example, spiking activity in chronically implanted electrodes may degrade over time faster than field potential modalities such as LFP or ECoG [41, 44, 58, 78]. Thus, combining spikes with field potentials can mitigate the impact of such degradation. Indeed, in our simulations, we find that multiscale SID improves the identification of both the shared modes and the distinct modes (Figure 5, A1).

### 3.6. Applications and future research directions

Given its computational efficiency, the multiscale SID method can enable various future real-time learning applications in neuroscience and neural engineering. For example, future work can utilize the multiscale SID method to develop an adaptive learning algorithm for tracking plasticity and non-stationarities in multimodal neural signals [22,25]. Adaptive tracking of neural dynamics is important for studying plasticity in the brain [79, 80] and for neurotechnologies that need to operate over long time periods, such as closed-loop deep brain stimulation (DBS) systems [81–87] or BMIs [81, 88–94]. In these technologies, various factors such as recording instability [95–97], learning and plasticity [58, 98–110], or a change in an internal state such as a psychiatric state [111] can lead to changes in multiscale dynamics. We recently developed an SID-based adaptive learning algorithm for single-scale continuous neural activity [25] and demonstrated its success in tracking of non-stationarity in multi-day ECoG recordings from epilepsy patients [22]. Developing a multiscale adaptive learning algorithm will be an important direction for future investigation.

Recent work have shown the benefit of learning a dynamical model for neural-behavioral data together by developing an SID-based method for two signal sources termed preferential SID or PSID [23]. Compared to modeling of neural dynamics unsupervised with respect to behavior, PSID preferentially learns the behaviorally relevant neural dynamics, thus achieving better neural decoding of behavior using lower-dimensional latent states [23]. But PSID models a single modality of neural activity so far. Thus, extending the multiscale SID method to consider multimodal neural data together with behavioral data during learning could be an interesting future direction. This could allow us to preferentially learn the multimodal neural dynamics that are behaviorally relevant and dissociate them from behaviorally irrelevant dynamics, thus potentially learning the former more accurately.

## 4. Methods

In this section, we provide the details for all analyses in simulations and in the non-human primate (NHP) data, as well as the detailed derivation of the multiscale SID algorithm. For a brief overview of multiscale SID please see section 2.1 and Algorithm 1).

### 4.1. Formulation of the multiscale dynamical model

We model the discrete-continuous spiking and field potential activity jointly as follows:

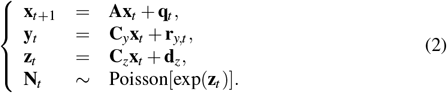

In this model, which we refer to as the multiscale dynamical model, we write a state-space model with latent states 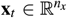, to jointly describe the dynamics of continuous Gaussian signals denoted by 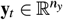 (e.g., field potentials) and discrete spike counts denoted by 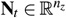. The temporal structure, i.e., dynamics, of the latent state **x**_*t*_ is described using an auto-regressive linear state equation with the state transition matrix 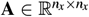. The continuous Gaussian signals **y**_*t*_ are modeled as a linear function of the latent states **x**_*t*_, and the discrete spike counts **N**_*t*_ are modeled as Poisson-distributed with latent log firing rate 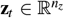 [43, 48, 63, 67, 76, 112, 113], which is a linear function of the same latent states. 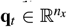 and 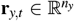 represent state and continuous observation noises and are modeled as uncorrelated white Gaussian noises with covariances 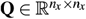 and 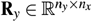, respectively. 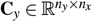 and 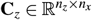 relate the continuous signal and the log firing rates to the latent state and **d**_*z*_ specifies the intercept (bias) of the log firing rate. The goal of multiscale SID is to learn all the multiscale model parameters *𝒩* = {**A, C**_*y*_, **C**_*z*_, **Q, R**_*y*_, **d**_*z*_} from multimodal neural training data.

In addition, we need to account for the potentially different time scales of observed spike counts and continuous signals such as field potentials. Indeed, we may observe spiking activity at every time step but have new samples of field potential features computed at every *M* ≥1 time steps. To account for such differences in the sampling rate of spike and field potential features, we model the field potential observations as missing observations for all the intermediate time steps where only spiking activity is observed but field potentials features are not computed.

The multiscale dynamical model formulation used here is similar to that for developing the multiscale EM algorithm [48, 49]. It is a generalized linear model (GLM) and is amenable to efficient and tractable inference, e.g., with the multiscale filter [43]. In prior work deriving and using the multiscale filter [43, 48, 49], spikes were modeled as a binary point process, where there is either 0 or 1 spike in each time step (1 ≥ *N*_*t*_ ≥ 0). Nevertheless, the derivation of the multiscale filter also holds for the case of a Poisson process, where there can be 0, 1 or more spikes in each time step (*N*_*t*_ ≥ 0), which we will use in this work.

It will be useful for our derivation of multiscale SID to rewrite the model from equation 2 in the following more general yet more compact form:

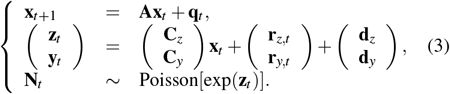

where 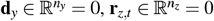 and

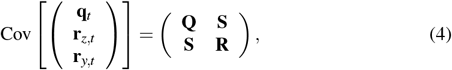

with

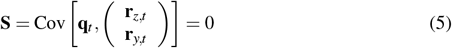

and

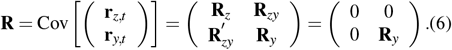

### 4.2. Multiscale SID algorithm

In this section, we first review the traditional covariance-based SID algorithm used for modeling single-modal Gaussian observations [61, 62], such as field potentials (Figure 1(a)). We then present the new multiscale SID algorithm for learning the multiscale dynamical model (Figure 1(b)).

#### 4.2.1. The single-scale SID algorithm for Gaussian observations

We model the single-modal Gaussian observations, e.g., field potentials, using the first two lines of equation 2 that describe the temporal evolution of the latent state and the Gaussian observation as:

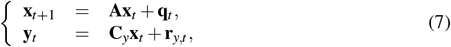

Briefly, the covariance-based SID algorithm finds the single-scale model parameters ℳ = {**A, C**_*y*_, **Q, R**_*y*_} as follows [62] (Figure 1(a)):

i. Form the future and past observation vectors 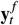 and 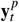 by stacking time-lagged observations as:

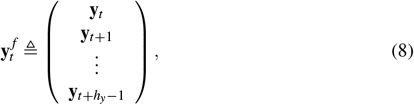

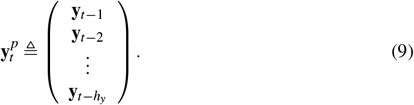

Here, *h*_*y*_ is the horizon hyper-parameter of the SID algorithm, which needs to be specified by the user manually. The horizon *h*_*y*_ must be larger than 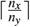 [61], such that the extended observability matrix 𝒪_*y*_ (equation 12) obtained in the next step can have full column rank. This is a necessary condition for the final learned state-space model to be observable, meaning that the latent states can be uniquely estimated from the observations.
ii. Empirically compute the future-past cross-covariance matrix **H**_*y*_ from the data formed in step i as 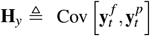. It is easy to see that **H**_*y*_ can be written in terms of auto covariances of **y**_*t*_ at different time delays *k*, i.e. in terms of 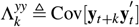, as:

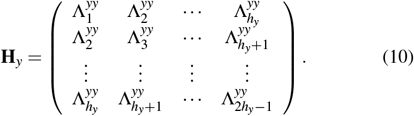

Since 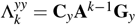 with 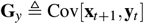 (see Appendix C), **H**_*y*_ can be decomposed in terms of **A, C**_*y*_ and **G**_*y*_, as:

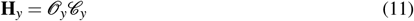

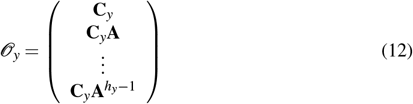

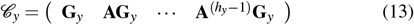

where 𝒪_*y*_ and 𝒢_*y*_ are termed extended observability matrix and extended reachability matrix, respectively [61, 62].
iii. Take singular value decomposition (SVD) of empirically estimated **H**_*y*_ from step ii to decompose it into the extended observability (𝒪_*y*_) and reachability (𝒢_*y*_) matrices (see [61, 62] for details). As shown in the previous step, these estimated 𝒪_*y*_ and 𝒢_*y*_ matrices will be functions of the yet-unknown model parameters **A, C**_*y*_ and **G**_*y*_.
iv. Find model parameters ℳ = {**A, C**_*y*_, **Q, R**_*y*_} from estimates of 𝒪_*y*_ and 𝒢_*y*_ and by solving a linear least squares problem and an algebraic Riccati equation. Interested readers can refer to [61, 62] for more details.

These steps conclude the traditional covariance-based SID algorithm for learning single-scale models with Gaussian observations.

#### 4.2.2. Outstanding challenges for developing a multiscale SID

The traditional covariance-based SID (reviewed in section 4.2.1) is not applicable to the multiscale model from equation 2 due to three standing challenges. In this section, we will explain these challenges for developing a multiscale SID and provide a brief explanation of our approach for addressing them in this section.

The first challenge in developing multiscale SID is that for the spiking modality, the log firing rates denoted by **z**_*t*_ are not observed. Rather, only a stochastic Poisson-distributed spike count time series, denoted by **N**_*t*_, is observed, which is nonlinearly related to the log-firing rates (equation 2). Thus, one cannot directly compute the empirical statistics of log firing rates as is possible with the Gaussian continuous modality **y**_*t*_. For example, prior work for single-modal spiking activity has computed the covariance of log firing rates by transforming the moments of the observable spike counts [63]. In our multiscale dynamical model, in addition to the covariances for each modality on its own, the cross-covariance terms between the two modalities are required for parameter learning. However, since spiking activity is one of the modalities, we cannot directly estimate the cross-covariance of its log firing rates with the continuous modality (**y**_*t*_, e.g., field potentials). To infer these cross-covariance terms, we use the method of moment transformation similar to prior work on single-modal spiking activity [63], but this time we find the transformation for the cross-covariance between discrete and continuous multimodal observations (section 4.2.3). Note that this prior work did not address the modeling of multimodal Poisson observations simultaneously with Gaussian observations—which also necessitates modeling of their joint statistics (equation 19)—, nor did it address the other two standing challenges in multiscale SID, that we outline next.

The second challenge in developing multiscale SID is related to ensuring that learned noise statistics are valid and conform to the multiscale model structure in equation set 3 and the assumptions of its latent state inference algorithm, i.e., the multiscale filter [43, 48]. Covariance-based SID methods [62], including the single-scale SID with Poisson observations [63], do not guarantee the validity of their learned noise covariance parameters. For example, these methods may learn noise covariance matrices **Q** or **R**_*y*_ that are not positive semidefinite (PSD), which is a necessary condition for a model representing real-valued time series and for enabling the statistical inference of its latent states. We address this challenge by devising a convex constrained optimization problem that finds a valid set of noise statistics for the model (section 4.2.6).

The third challenge in developing multiscale SID is that Gaussian continuous observations such as field potentials **y**_*t*_ and spike count observations **N**_*t*_ may be available at different timescales due to differences in their sampling rate. This means that some timesteps may have the slower modality as missing observations. We address this challenge by resampling and interpolating the slower signals within the training data and prior to forming the noise covariances, which can work when the Nyquist sampling rate criterion is met for the slower signal as is often the case for field potential signals (section 4.2.3). Once model parameters are learned, we no longer need to perform interpolation in the test set; instead, we will estimate the latent states and predict the neural activity using the multiscale filter (MSF) [43], which can process multimodal data with different sampling rates.

In the following sections, we provide details of how we address each of these challenges and finally estimate all the model parameters *𝒩* = {**A, C**_*y*_, **C**_*z*_, **Q, R**_*y*_, **d**_*z*_}.

#### 4.2.3. Empirical estimation of the future-past cross-covariances between the log firing rates and the continuous modality (i.e., **H**_w_)

To model multimodal data per equation 2 (Figure 1(b)), we first empirically compute the following future-past cross-covariance matrix **H**_*w*_ and subsequently estimate model parameters from it:

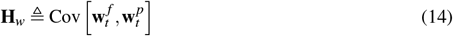

with

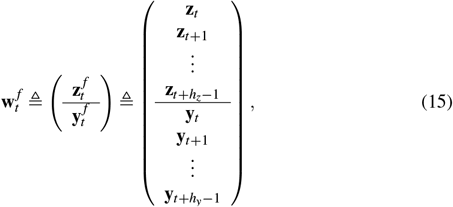

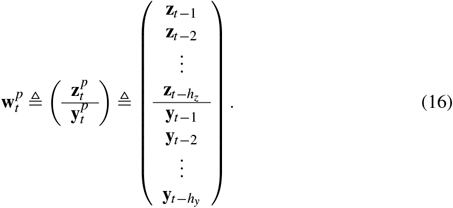

Here 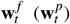 is formed by stacking the future (past) latent log firing rate vector and the future (past) observed continuous modality vector. *h*_*z*_ is another horizon parameter that needs to be selected manually similar to *h*_*y*_ (section 4.2.1) and corresponds to spiking activity. In all analyses in this work, we set horizon parameters as *h*_*y*_ = *h*_*z*_ = 10 unless otherwise stated.

Note that log firing rates **z**_*t*_ are not directly observed, and thus **H**_*w*_ (equations 14-16) cannot be directly estimated from data as a sample covariance. To estimate **H**_*w*_ from data, we need estimates of auto covariances of **y**_*t*_ as well as auto covariances of **z**_*t*_ and cross-covariances of **y**_*t*_ and **z**_*t*_ at different time delays. While we can directly estimate moments of **y**_*t*_ from the continuous observations, we cannot do the same for auto covariances of **z**_*t*_ and cross-covariances of **y**_*t*_ and **z**_*t*_ as log firing rate **z**_*t*_ is not observable. However, we can use the fact that the model in 2 dictates a computable relationship between moments of **y**_*t*_ and **z**_*t*_ and those of **y**_*t*_ and **N**_*t*_, which can be directly estimated from the discrete-continuous observations. So we transform moments [63] that are directly computable from data (i.e., moments of **y**_*t*_ and **N**_*t*_ and their cross-terms) to the unknown moments required to estimate **H**_*w*_, i.e., moments of **y**_*t*_ and **z**_*t*_ and their cross-terms, i.e., the cross-terms between the two modalities.

To do so, first we define the future-past vector of the continuous modality activity 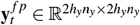 by stacking 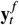 and 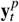 as:

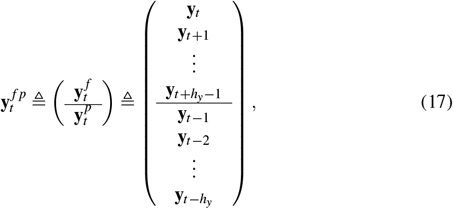

Similarly we define 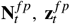 for variables **N**_*t*_ and **z**_*t*_ by stacking the corresponding future and past vectors. Then we define the mean denoted by *μ*, and the auto-covariance and cross-covariance denoted by Σ of these variables as follows:

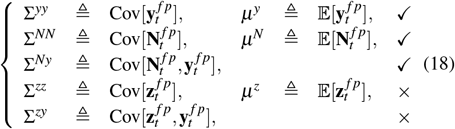

We have empirical estimates of Σ^*yy*^ and *μ*^*y*^ directly from the continuous observations, Σ^*NN*^ and *μ*^*N*^ directly from spike observations and Σ^*Ny*^ directly from both observations (first three lines in equation set 18). These empirical moments correspond to the output of moment computation block in Figure 1(b). We then compute Σ^*zy*^, Σ^*zz*^ and *μ*^*z*^ (last two lines in equation set 18) that are not directly computable from observations, by a moment transformation procedure based on the following relations:

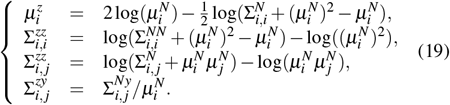

where ._*i*_ refers to the *i*th element of a vector, and ._*i, j*_ to the element in the *i*th row and *j*th column of a matrix. In Appendix A, we derive the relation for computing Σ^*zy*^ in equation set 19, which is the cross-covariance between the two discrete-continuous modalities. The relations for computing *μ*^*z*^ and Σ^*zz*^ in equation set 19 are derived in [63] where single-scale SID is derived for a single modality with Poisson distribution.

The above procedure addresses the first challenge in developing multiscale SID (see section 4.2.2), giving an estimate of the future-past cross-covariance matrix **H**_*w*_ (Figure 1(b)). See Appendix B for constructing **H**_*w*_ based on equations 18 and 19. We also compute 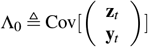 from the quantities computed in equation 18 (Appendix B), to be used when we subsequently estimate all the model parameters *𝒩* = {**A, C**_*y*_, **C**_*z*_, **Q, R**_*y*_, **d**_*z*_} using the estimated **H**_*w*_, Λ_0_ and *μ*^*z*^ later in section 4.2.5.

To address the challenge of potentially different sampling rates in developing multiscale SID (see section 4.2.2), we proceed as follows. For empirical estimation of Σ^*yy*^ and Σ^*Ny*^ from the discrete-continuous observations in the training data, we first interpolate the slower modality to make the sampling rate of the two modalities the same. Here, we assume that we observe the spiking activity at every time step, while we may compute the continuous modality such as field potential power features every *M* ≥ 1 time steps (section 4.1) and therefore have continuous observations **y**_*t*_ every *M* ≥ 1 time steps. This is often the case in neural spike-field datasets [43, 49, 56]. Thus, we increase the sampling rate of the continuous observations by a factor of *M* by filling in their missing samples with zeros and then applying a zero-phase FIR filter (see [114], we use the “interp” command in MATLAB).

It is worth noting that by performing interpolation for the slower modality, we assume that the multiscale data is collected at an appropriate sampling rate for each modality, such that information has not already been irreversibly lost due to aliasing when the data was originally sampled [115] – i.e., we assume that Nyquist sampling rate requirements are met. This assumption is reasonable because any information lost due to aliasing is not retrievable by any learning method and thus interpolation to recover the existing information is a reasonable approach. Moreover, note that this interpolation is only needed in the training data for computations of equation 18 when learning the model parameters and not for prediction of neural activity or behavior after the model parameters are learned.

We address the challenge of ensuring valid noise statistics in sections 4.2.5 and 4.2.6 after first presenting how model parameters relate to the computed covariances.

#### 4.2.4. Relation of **H**_w_ to the model parameters

In this section, we will show how **H**_*w*_ (equation 14) can be written in terms of the multiscale model parameters, which we will later use to extract the model parameters from **H**_*w*_ in section 4.2.5. As discussed in section 4.2.3, we can write **H**_*w*_ in terms of the cross and auto covariances of **y**_*t*_ and **z**_*t*_ at different time delays *k*, i.e. in terms of 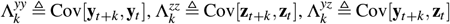 and 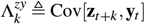 as:

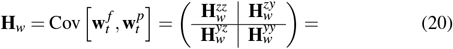

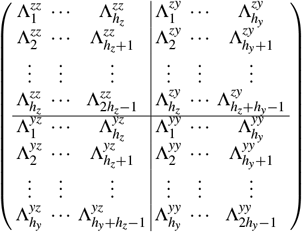

We can then write **H**_*w*_ in terms of model parameters **A, C**_*y*_, **C**_*z*_, **G**_*y*_, **G**_*z*_, where **G**_*y*_ = Cov[**x**_*t*+1_, **y**_*t*_] and **G**_*z*_ = Cov[**x**_*t*+1_, **z**_*t*_]. It can be shown (see Appendix C) that for positive integer *k*’s we have 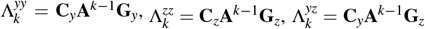 and 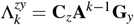. Replacing these values in equation 20, gives:

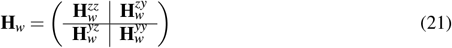

With

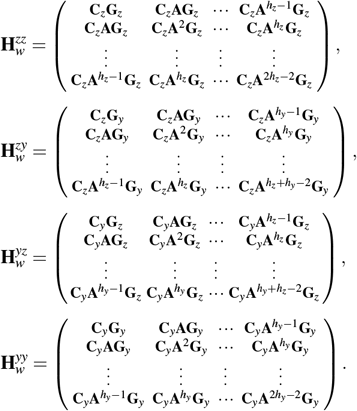

It is easy to see from equation 21 that **H**_*w*_ can be decomposed as:

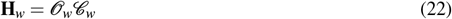

with

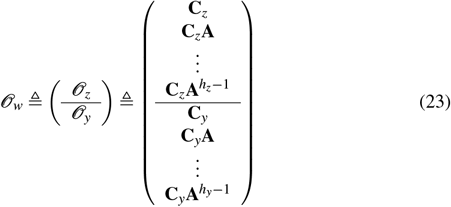

and

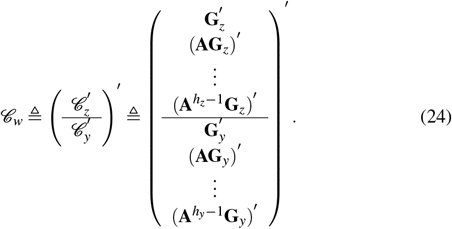

Here 𝒪_*w*_ (𝒞_*w*_) is the multiscale extended observability (reachability) matrix, which is the concatenation of single-scale extended observability (reachability) matrices, 𝒪_*z*_ and 𝒪_*y*_ (𝒞_*z*_ and 𝒞_*y*_).

This concludes how **H**_*w*_ is related to model parameters. Based on these relationships, we will next use the estimation of **H**_*w*_ obtained from real data (see section 4.2.3) to estimate all model parameters.

#### 4.2.5. Estimating model parameters from empirical estimates of **H**_w_, Λ_0_ and μ^z^

Using the the following steps, we estimate the multiscale model parameters *𝒩* = {**A, C**_*y*_, **C**_*z*_, **Q, R**_*y*_, **d**_*z*_} from the estimated **H**_*w*_ and Λ_0_ and *μ*^*z*^, which were estimated from the data via the transformation of moments technique (section 4.2.3):

i. Find estimates of extended observability matrix 𝒪_*w*_ and extended reachability matrix 𝒞_*w*_ (equations 22-24) by applying SVD on the estimated **H**_*w*_ and keeping only the largest *n*_*x*_ singular values:

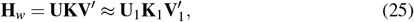

Here 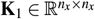 is a diagonal matrix containing the *n*_*x*_ largest singular values, and 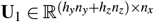 and 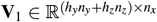 are the associated left and right singular vectors, respectively. We then have:

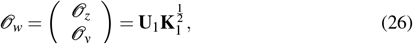

and

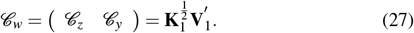
ii. Extract estimates of **C**_*y*_ and **C**_*z*_ as the first *n*_*y*_ and *n*_*z*_ rows of estimates of 𝒞_*y*_ and 𝒞_*z*_ from step (i), respectively (see equation 23):

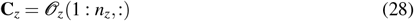

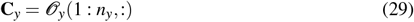

where : is used to indicate selecting all elements along a given row or column. Further *m* : *n* indicates selection of elements ranging from the *m*-th to the *n*-th position along a row or column.
iii. Estimate **A** by solving the following optimization problem:

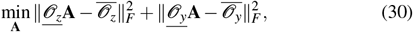

Where

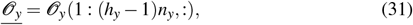

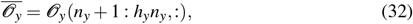

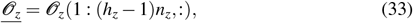

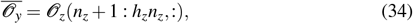

and ∥.∥_*F*_ represents the Frobenius norm. The optimization problem in 30 combines information across modalities, i.e. multimodal discrete-continuous data, through the cost function which sums up squared error of finding **A** from 𝒪_*z*_ and from 𝒪_*y*_ (see equation 23). The optimization problem in 30 can also be written as:

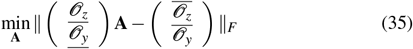

which has the following analytical least square solution:

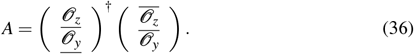
iv. Extract **G**_*z*_ and **G**_*y*_ as the first *n*_*y*_ and *n*_*z*_ columns of estimates of 𝒞_*z*_ and 𝒞_*y*_ from step (i) (see equation 24): **G**_*z*_ = *C*_*z*_(:, 1 : *n*_*z*_), (37)

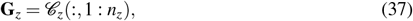

and

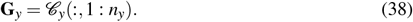
v. Estimate valid noise covariances **Q** and **R**_*y*_ by solving the following convex constrained optimization problem, which addresses the second challenge in developing multiscale SID (section 4.2.2):

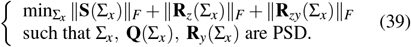

where

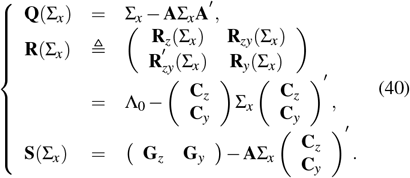

With 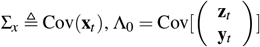. Note that the estimates for all the model parameters that are required to solve this optimization problem – i.e., **A, C**_*y*_, **C**_*z*_, **G**_*y*_, **G**_*z*_, Λ_0_ – are available from previous steps. Also, equations in 40, can be derived from the model in equation 3 (see Appendix D). For more details and a description of how this step (along with step vi) addresses the second challenge in developing multiscale SID see section 4.2.6.
vi. Update estimates of **G**_*y*_, **G**_*z*_, Λ_0_: given the solution for Σ_*x*_ obtained from solving the constrained optimization in the previous step, we set **R**_*z*_, **R**_*zy*_ and **S** to exactly 0 in equation set 40, and get updated estimates for Λ_0_, **G**_*y*_ and **G**_*z*_. (vii) Read **d**_*z*_ from the estimated first moment of future-past log firing rate vector, i.e. 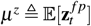 in section 4.2.3:

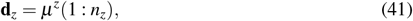

This concludes the learning of all multiscale model parameters *𝒩* = {**A, C**_*y*_, **C**_*z*_, **Q, R**_*y*_, **d**_*z*_}. All steps of multiscale SID algorithm are summarized in Algorithm 2.

Finally, we note that 𝒫 = {**A, C**_*z*_, **C**_*y*_, **G**_*z*_, **G**_*y*_, Λ_0_, **d**_*z*_, Σ_*x*_} is also an alternative full specification of the multiscale model, which is equivalent to the specification with *𝒩* = {**A, C**_*y*_, **C**_*z*_, **Q, R**_*y*_, **d**_*z*_}. This is due to the one to one relation between these two sets according to equations in 40 [23]. The model specification with *𝒩* is useful when using the multiscale filter for neural or behavior prediction (sections 4.3.3,4.4.3) [43], while 𝒫 is more useful for model parameter evaluation (section 4.3.2).

#### 4.2.6. Ensuring validity of noise statistics

In steps v-vi of the parameter estimation procedure described in section 4.2.5, we addressed the key challenge of ensuring valid noise statistics in developing multiscale SID (section 4.2.2) by devising a convex constrained optimization. In this section, we provide more details and context for this novel approach. The optimization problem in equation 39 aims to find noise statistics that satisfy the following conditions that are required in the multiscale model (equation 3):

i. State and continuous observation noise covariances **Q** and **R**_*y*_ must be valid covariance matrices, and thus need to be positive semidefinite.
ii. State and continuous observation noises are assumed to be uncorrelated, i.e. **S** = 0.
iii. Log-firing rate is assumed not to have additive Gaussian noise, i.e., **R**_*z*_ = 0 [43, 49] with the stochasticity of spiking data reflected in its Poisson distribution.

The last two conditions are incorporated in the multiscale model (equation 3) to be consistent with prior work on Poisson and multiscale model structures [43, 49, 63, 76], but technically could be relaxed if a future extension of the multiscale model/filter drops those assumptions [43, 49]. To encourage solutions that satisfy the above conditions as closely as possible, in the optimization problem of equation 39, we minimize the sum of norms of **S**(Σ_*x*_), **R**_*z*_(Σ_*x*_) and **R**_*zy*_(Σ_*x*_), subject to the required condition that Σ_*x*_, **Q**(Σ_*x*_) and **R**_*y*_(Σ_*x*_) are PSD. We find the numerical solution of Σ_*x*_ in the convex constrained optimization problem in 39 [71, 72] using CVX [70], a MATLAB software that uses a disciplined convex programming approach [116]. We then get estimates of **Q, R**_*y*_ according to equation set 40. Alternatively, one could form and solve a similar constrained optimization problem using generic numerical optimization tools rather than using semi-definite programming via CVX [70]; the former may be less accurate but useful if equation 40 gives an infeasible convex optimization problem. This might happen if **H**_*w*_ and subsequently other parameters before noise statistics estimation are poorly estimated due to short length of data or low signal-to-noise-ratio (SNR).

##### Algorithm 2 Multiscale SID

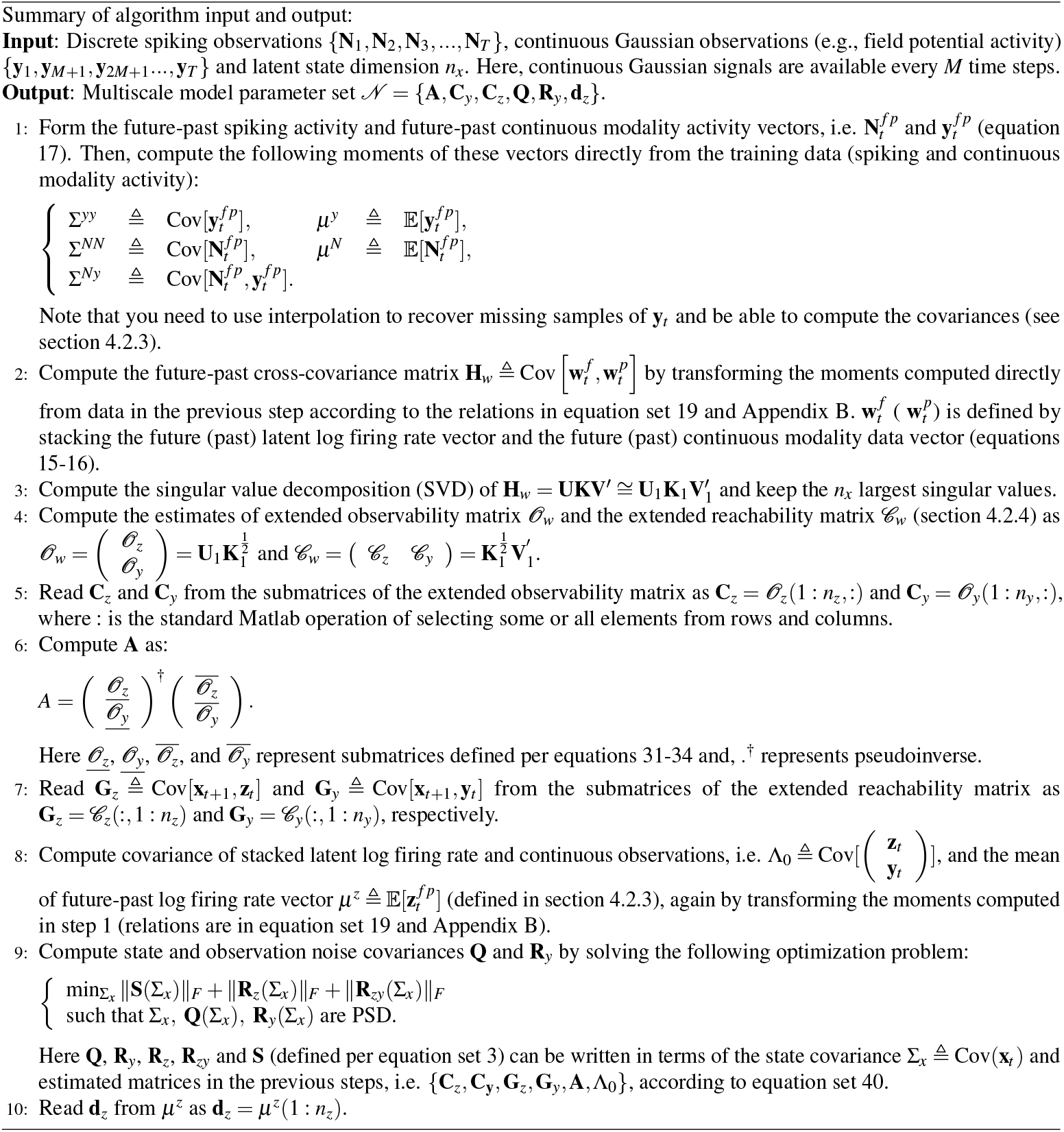

The fundamental reason why we have the flexibility to find alternative sets of noise statistics that satisfy conditions such as the above three is that the states in the multiscale model (equation 3) are latent and thus there are infinitely many equivalent solutions with different latent states for describing the same observed multimodal data **y**_*t*_ and **N**_*t*_. These include, but are not limited to (see Faurre’s theorem in [61, 62]), all equivalent models obtained by left-multiplying the latent state with an arbitrary invertible matrix, also known as similarity transformations. These equivalent alternative models have different state covariance matrices Σ_*x*_. The optimization problem in equation 39 aims to find one of these equivalent models that satisfies the required conditions as much as possible. This approach is very flexible and thus could be useful for other problems, for example to guarantee validity of noise statistics in all variants of covariance-based SID in general whether single-scale or multiscale (see Discussion).

### 4.3. Validation of multiscale SID using simulations

#### 4.3.1. Simulating multimodal neural data

To validate our multiscale SID in numerical simulation, we randomly generate sets of multiscale model parameters *𝒩* ={**A, C**_*y*_, **C**_*z*_, **Q, R**_*y*_, **d**_*z*_} in equation 2 and then generate multimodal spike-field activity from these models. In random generation of the model parameters, we also set criteria for desired signal to noise ratio (SNR) of field features, bias and maximum firing rates of spikes, contribution of dynamical modes in each modality and range of frequency and magnitudes from which dynamical modes are randomly drawn, all of which will be explained later. Note that each dynamical mode corresponds to a pair of complex conjugate eigenvalues or a real eigenvalue of the state transition matrix **A** (Figure 1(d)).

Prior work suggests that there exist both shared and distinct dynamical modes in spiking and field potential activity [49]. Motivated by this and also to cover a general scenario, we simulate both shared and distinct modes. Distinct spike (field) modes are those that are only present in spiking (field) activity and shared modes are those that are present in both modalities of neural activity (both spiking and field potential activity). To quantify the presence of a mode in the dynamics of a modality, we define the contribution of the dynamical mode *i* to the dynamics of a modality cntrb_modality,*i*_ as the total variance of the activity in that modality that is generated from that mode (across all neural dimensions from that modality, either log firing rates **z**_*t*_ or field potential activity **y**_*t*_), i.e.:

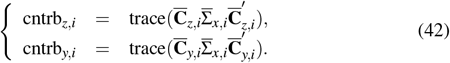

where 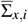 is the covariance of states corresponding to mode *i*, which is a submatrix of state covariance Σ_*x*_ (equation 54, Appendix E). Further, 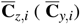 is a submatrix of **C**_*z*_ (**C**_*y*_) with columns corresponding to mode *i*. Note that, without loss of generality, **A** in simulation is generated in the block diagonal format (see item ii below). Finally, we denote the contribution of mode *i* to the dynamics of a modality normalized by the sum of contribution of all the modes as nCntrb_modality,*i*_.

To generate the multiscale model parameters *𝒩* = {**A, C**_*y*_, **C**_*z*_, **Q, R**_*y*_, **d**_*z*_} in equation 2, we proceed as follows:

i. Generate 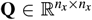: The diagonal entries of the diagonal **Q** matrix are absolute values of samples drawn from the standard normal distribution.
ii. Generate 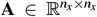: We generate dynamical modes as *re* ^*jθ*^, with randomly choosing *r* between [0.950, 0.999], and *θ* between [0, 0.0316], consistent with ranges observed in prior work modeling the motor cortical activity of NHPs [49]. *r* and *θ* determine the damping and oscillatory behavior of the dynamics. Complex modes appear as complex conjugate eigenvalues of **A**. We construct **A** in a block diagonal format where the block corresponding to mode *i* is 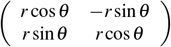. Note that the block diagonal construction of **A** does not pose any loss of generality since all models can be converted to this form (known as the canonical form) via a similarity transformation [117].
iii. Generate 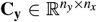 and 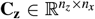 : Randomly generate entries of these matrices according to a uniform distribution. Then, scale columns and rows of these matrices to enforce the desired maximum firing rate for each neuron and the required contributions of each mode, cntrb_modality,*i*_ based on its type (see Appendix F for details). The maximum and bias of the firing rates are randomly and uniformly picked in the ranges [40Hz, 50Hz] and [5Hz, 10Hz], respectively.
iv. Generate 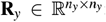: Having set **C**_*y*_ and Σ_*x*_ (Appendix E), we set diagonal entries of the diagonal **R**_*y*_ to achieve the desired SNRs for **y**_*t*_. The SNR vector of field potential features is defined as diag 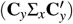.*/*diag(**R**_*y*_) and the entries are randomly picked in the range [0.8, 1.2]. Here, diag(.) denotes the operation of transforming the diagonal entries of a matrix into a vector, and “.*/*” is an element-wise division operator.

Given the multiscale model parameters *𝒩* = {**A, C**_*y*_, **C**_*z*_, **Q, R**_*y*_, **d**_*z*_} from equation 2, we can generate the multimodal spiking activity **N**_*t*_ for *t* ∈ {1, 2, 3…, *T*} and field potential activity **y**_*t*_ for *t* ∈ {1, *M* + 1, 2*M* + 1, …, *T*} as follows. We set **x**_**0**_ = 0 and generate **q**_*t*_ and **r**_*y,t*_ from zero-mean Gaussian distributions with covariance of **Q** and **R**_**y**_, respectively. We then generate **x**_*t*_, **y**_*t*_ and **z**_*t*_ by iterating through equation 2 for *t* = 1 to *t* = *T*. We set *M* = 5 here and discard field potentials at the intermittent time steps as missing observations. **N**_*t*_ is then generated from Poisson distributions with rates equal to the elements of the vector exp(**z**_*t*_).

#### 4.3.2. Quantifying parameter identification error

There are infinitely many equivalent latent state models for the multiscale model in equation 2, for example any invertible linear mapping of the latent state is a similarity transform that gives an equivalent model [23, 61, 62]. To take this into account when evaluating the learned models in simulations, we first find the similarity transform that makes the learned model as close as possible to the true model in simulations in terms of the basis of the latent state as also done in prior work [23] (see Appendix G for details). We then compare the model parameters for the transformed learned model with the true model parameters. Note that this procedure does *not* change the learned model, rather only gives a different equivalent formulation for it so that it can be compared with the true model [23].

Given the true and the learned model parameters, we quantify the parameter identification error of a matrix/vector parameter *ψ* as:

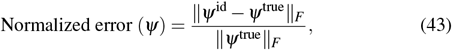

where ∥.∥ _*F*_ represents Frobenius norm and *ψ*^true^ and *ψ*^id^ refer to the true and identified parameter values, respectively. We evaluate our multiscale SID algorithm by computing the normalized error for each *ψ* ∈ 𝒫 except for Σ_*x*_. This is because according to Faurre’s theorem [23, 61, 62], all the model parameters in 𝒫 other than Σ_*x*_ are uniquely determined from **y**_*t*_ and **N**_*t*_ up to within a similarity transform. Σ_*x*_ on the other hand is a redundant description (or internal description, see [62]) for the observations and may have infinitely many solutions even beyond similarity transforms for the same observations [23, 62], and thus even a perfect learning method may not need to learn the same Σ_*x*_ (within a similarity transform) as the true model [23].

We also compute the normalized error for the vector of eigenvalues of **A**; i.e., *ψ* = eig(**A**), and denote it as normalized mode error (see Appendix H for more details). Note that the vector of eigenvalues of **A** does not change after similarity transformation up to within permutations. Furthermore, in addition to computing the normalized mode error for all the modes at once, we can compute it separately for each mode type – distinct spike modes, distinct field modes and shared modes (section 4.3.1) – to investigate how multiscale SID identifies each and the collective dynamics of both modalities.

For this analysis, we simulate 50 multiscale dynamical models according to section 4.3.1 with dimension of spiking activity *n*_*z*_ randomly picked in the interval [10, 30], and set dimension of field potential activity *n*_*y*_ = 4 × *n*_*z*_. We also set number of dynamical modes to four, with two shared modes, one distinct spike mode and one distinct field mode. To evaluate the effect of training sample size on these errors, we generate multimodal spiking and field potential activity with different sample sizes *T* ∈ [1, 2, 10, 50, 100, 1000] × 10^3^ from each model.

#### 4.3.3. One-step-ahead prediction of spiking and field potential activity

To obtain the one-step-ahead prediction of spiking and field potential activity, we need to obtain the one-step-ahead prediction of latent states as a first step. To do so, we use the identified model parameters to construct the optimal filters to obtain the one-step-ahead prediction of latent states (Figure 1). The optimal filters for the single-modal spiking activity, the single-modal field potential activity and the multimodal spiking and field potential activity are the point process filter (PPF), the Kalman filter (KF) and the multiscale filter (MSF), respectively. MSF is derived in our prior work and in special cases when only one of the two signals is observed, it reduces to either PPF or KF [43]. MSF can also simultaneously admit modalities that have different sampling rates by treating the intermediate samples of the slower modality as missing, thus not requiring interpolation [43]. We denote the one-step-ahead prediction of latent states at time step *t* as **x**_*t*|*t*−11_, where **O**_*t*|*t*−1_ denotes an estimation of **O**_*t*_ based on all neural observations up to time step *t* − 1.

Given the one-step-ahead prediction of latent states **x**_*t*|*t*−1_, the one-step-ahead prediction of field potentials is **y**_*t*|*t*−1_ = **C**_*y*_**x**_*t*|*t*−1_. We use Pearson’s correlation coefficient (CC) between the one-step-ahead predicted field potential activity **y**_*t*|*t*−1_ and the true field potential activity **y**_*t*_, averaged over dimensions of field potential activity *n*_*y*_, as the accuracy measure for the one-step-ahead prediction of field potential activity [49].

We also obtain the one-step-ahead prediction of log firing rate as **z**_*t*|*t*−1_ = **C**_*z*_**x**_*t*|*t*−1_ + **d**_*z*_. We then construct the receiver operating curve (ROC) for deciding which time steps (bins) contained at least one spiking event [6, 47, 49]. We then use prediction power defined as PP = 2AUC−1, averaged over spiking dimensions *n*_*z*_, as the accuracy measure for the one-step-ahead prediction of spiking activity. Here AUC stands for area under curve of the ROC.

In these set of simulations we compute one-step-ahead prediction of spiking and field potential activity on a test set with 10^4^ samples, using the identified model parameters from the training set.

#### 4.3.4. Comparison of multiscale SID and multiscale EM in computation cost and accuracy

We compare the multiscale SID with the multiscale EM, which is the current method for learning the multiscale dynamical model [48, 49]. We perform comparisons in terms of computation cost and accuracy in identifying the dynamics and in one-step-ahead prediction of spiking and field potential activity.

Multiscale EM is an iterative algorithm [48, 49], which similar to other EM methods iteratively maximizes the data likelihood [76, 118, 119] but this time for multiscale discrete-continuous data. We continue the iterations until the following convergence criterion is met or until the number of iterations has reached a predefined maximum allowable number of iterations, which is set to 175 iterations here. We set the convergence criterion of the multiscale EM by putting a threshold on the relative change of a performance measure *m* in two consecutive steps, i.e.:

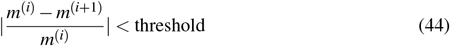

where *m*^(*i*)^ represents the performance measure *m* at iteration *i*. In this simulation analysis we take the normalized mode error (see section 4.3.2) as the performance measure *m* and we set the threshold to 10^−4^.

To compare the computation costs of multiscale SID vs. multiscale EM, we report the time it takes to learn the model parameters by each of the algorithms on the same computer. Further, to compare the performance of these algorithms in terms of identification of dynamics, we report the normalized mode error (section 4.3.2). Finally, we compare the accuracy in one-step-ahead prediction of spiking and field potential activities, which are quantified by PP and CC, respectively (section 4.3.3). These variables are reported for different training sample sizes for the same 50 simulated multiscale models as in section 4.3.2.

#### 4.3.5. Comparison of multiscale SID and single-scale SID in identification of dynamics

To demonstrate the potential benefit of the multiscale modeling over single-scale modeling for identification of dynamics, we perform an analysis where we add neural signal of one modality to the other modality in simulated data.

We first simulate 50 multiscale dynamical models (equation 2) with random parameters according to section 4.3.1, with *n*_*z*_ = 14 spiking signals, *n*_*y*_ = 14 field potential signal, and the same number of shared and distinct modes as previous sections. We generate multimodal spiking and field potential activity with *T* = 10^5^ samples from each model. Then, we construct and model sub-networks of the simulated multimodal network activity by gradually adding more signals from one modality (either spiking or field potential activity, *n*_add_ ∈ {2, 4, …, 14}), to a fixed number of neural signals from the other modality, denoted as baseline neural signals. We set the number of baseline neural signals *n*_baseline_ = 4, 6 or 14. We study how the learning error for the dynamics, quantified by normalized mode error (see section 4.3.2), changes as signals from one modality of neural activity are added to signals from the other modality. In addition, we study how this error changes for each mode type – i.e., distinct spiking or field modes versus shared modes – separately (sections 4.3.1 and 4.3.2). With this we demonstrate how multiscale modeling helps to model the collective dynamics of both modalities, which includes both the modes that are only present in one of the two modalities and the shared modes that are present in both modalities. In this analysis, when both modalities are observed, we use our multiscale SID; when only baseline field potential or only baseline spiking activity is observed, we use the traditional single-scale SID algorithm for Gaussian observations [61] or the single-scale SID algorithm for Poisson observations [63], respectively.

### 4.4. Validation of multiscale SID using NHP dataset

#### 4.4.1. Neural and behavioral recordings

We model the neural and behavioral data recorded from a male non-human primate (NHP) (Monkey J), as it performed naturalistic 3D reach and grasp movements for a liquid reward (Figure 6(a), see [23, 49, 69] for more details). All surgical and experimental procedures were in compliance with National Institute of Health Guide for Care and Use of Laboratory Animals and were approved by New York University Institutional Animal Care and Use Committee. A 137 electrode microdrive (Gray Matter Research, USA) was used to record spiking and local field potential (LFP) activity from left hemisphere motor cortical areas, covering parts of the primary motor cortex (M1), the dorsal premotor cortex (PMd), the ventral premotor cortex (PMv), and the prefrontal cortex (PFC). Angle of multiple joints in the active arm (right) were inferred from the tracked position of retroreflective markers placed on the arm by using a NHP musculoskeletal model and inverse kinematics (SIMM, MusculoGraphics Inc., USA) [120]. We predict the angle of the following 7 prominent joints in our analyses: shoulder elevation, elevation angle, shoulder rotation, elbow flexion, pro supination, wrist flexion, and wrist deviation [49].

#### 4.4.2. Neural data processing

To obtain the spiking activity (**N**_*t*_), spiking events were detected every time the band pass filtered raw neural signals (filtered within 0.3-6.6kHz) crossed a threshold of 3.5 standard deviations below their mean [69], and were counted in 10ms bins to get **N**_*t*_. To obtain LFP features (**y**_*t*_), we first low pass filtered the raw neural signals with cut off frequency of 400 Hz and then down sampled it to 1 kHz. We then for each channel computed log-powers in seven frequency bands: theta (4–8 Hz), alpha (8–12 Hz), beta 1 (12–24 Hz), beta 2 (24–34 Hz), gamma 1 (34–55 Hz), gamma 2 (65–95 Hz), and gamma 3 (130–170 Hz) [43, 49, 59]. The log-power features were computed by first performing common average referencing, and then computing short-time Fourier transform for causal sliding windows of 300 ms every 50 ms. Thus, for our analyses, the time scale of LFP log-power features was 50 ms, and that of spike events was 10 ms [43, 49].

#### 4.4.3. Predicting behavior from the estimated latent states

To predict the behavior, i.e., joint angle trajectories, from the NHP neural data (section 4.4.1), we first use the learned models to estimate the low-dimensional latent states **x**_*t*|*t*_ from neural data, and then build a linear regression from these latent states to the behavior in the training set. To estimate the latent states **x**_*t*|*t*_, we use the learned model parameters to construct the associated optimal filters depending on the modality of neural activity observed in the model, i.e., KF, PPF or MSF [43](Figure 1(c)).

To build the regression model that predicts the behavior from latent states, we estimate the latent states within the training data and then compute the projection matrix **L**, which minimizes the mean squared error of predicting the behavior in the training data as 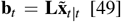 [49]. Here, 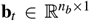 is the behavior where *n*_*b*_ = 7 denotes its dimension, and 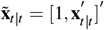 is the estimated latent state vector concatenated with a constant to account for bias. The solution to this for this ordinary least squares linear regression problem is:

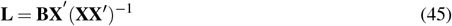

where, 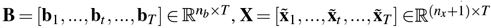 and *T* is the size of training set. In the test set, we first estimate the latent states **x**_*t*|*t*_ using the appropriate filter (MSF, KF, or PPF), and then predict the behavior using the learned **L** projection matrix from the training set (Figure 1(c)):

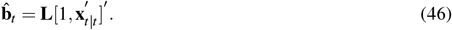

We use Pearson’s correlation coefficient (CC) between the predicted behavior and the true behavior, as the measure of behavior prediction accuracy. In our analysis, we report the mean of this correlation coefficient over the seven joint angle trajectories.

#### 4.4.4. Five-fold cross-validation

In all the analyses for the NHP dataset, we use five-fold cross-validation. More precisely, we divide the data from each experimental session into five equal sized continuous sections, and in each fold, use four out of the five sections for training and use the remaining section for testing. We repeat this procedure five times so that each section has been used as the test data once. Further, we perform our analyses across seven experimental sessions from the subject [23, 49]. Finally, in each cross-validation fold, we z-score each dimension of the field potential activity based on its mean and variance within the training set [23]. This was done as a preemptive measure to ensure that learning methods do not discount any dimension of the field potential activity even if that dimension had a much smaller variance compared with other dimensions [23].

#### 4.4.5. Comparison of multiscale SID and multiscale EM in terms of computation cost and accuracy

We predict neural activity and behavior, i.e., the seven joint angle trajectories (section 4.4.1), from the NHP multimodal spiking and field potential activity using both multiscale SID and EM and compare these algorithms in terms of accuracy and computation cost. For this analysis, we construct the multimodal spiking and field potential activity that is to be modeled for each recording session by picking the top 15 spike channels and the top 15 LFP channels with highest behavior prediction accuracy when modeled as individual channels. The behavior prediction accuracy of the individual channels is computed and sorted using a basic non-latent Kalman filter decoder where the states are taken to be the behavior itself [43, 121]. We identify the multiscale model parameters by applying the multiscale SID (section 4.1) or the multiscale EM [48, 49] to the training data. For each learned model, we then estimate the latent states in the test data using the MSF associated with the learned model, and predict the neural activity (section 4.3.3) and the behavior (section 4.4.3) from the estimated latent states (Figure 1(c)). We repeat this analysis for latent state dimensions *n*_*x*_ ∈ {2, 4, …, 24}. As with all other analyses, we set the horizon for multiscale SID as *h*_*y*_ = *h*_*z*_ = 10 (section 4.2.3). Finally, across different latent states, we compare the computation time it takes to learn the model parameters (similar to section 4.3.4), the one-step-ahead prediction accuracy of spiking and field potential activity, and the behavior prediction accuracy (defined in sections 4.3.3 and 4.4.3) between multiscale SID and multiscale EM algorithms. To determine the multiscale EM convergence, we set the measure *m* in equation 44 once to one-step-ahead prediction accuracy of field potential activity (CC) and once to that of spiking activity (PP) and take the larger *i* across the two as the convergence iteration. Similar to section 4.3.4, the multiscale EM is terminated when the convergence criterion is met or once we reach the predefined maximum allowable number of iterations, which we set to 150 iterations for this analysis.

#### 4.4.6. Comparison of multiscale SID and single-scale SID in predicting behavior

To investigate the potential benefit of multiscale modeling over single-scale modeling in the NHP dataset, we do a neural signal addition analysis similar to what we do for simulated data (section 4.3.4). We then study the behavior prediction accuracy instead of identification of dynamical modes as the ground-truth of the latter is not known in real data.

In this analysis, we pick the top 30 spike channels and the top 30 LFP channels (210 LFP power features), which have the highest single channel behavior prediction accuracy when modeled individually, as the spiking and field potential activity to be modeled. We then randomly select *n*_baseline_ signals from one modality, denoted as baseline neural signals, and gradually add additional randomly selected signals from the other modality of neural activity to them (in steps of *n*_add_ ∈ {2, 4, …, 14}). We repeat this process of random selection of baseline and added neural signals 10 times for each of *n*_baseline_ = 2, 6 or 14. For each pair of baseline neural signals and added neural signals, we use the multiscale SID in combination with the MSF to estimate the latent states, predict the behavior and compute the behavior prediction accuracy, all within a five-fold cross-validation (Figure 1(b)-(c), section 4.4.4). When evaluating models of baseline neural signals alone (not combined with the other modality of neural activity), we use the appropriate single-scale SID and single-scale filters to obtain the behavior prediction and compute their cross-validated accuracy (section 4.4.3). Given the computed behavior prediction accuracies for single-modal baseline neural signals, and for multimodal baseline and added neural signals together, we can study how multiscale modeling and filtering may help in behavior prediction compared to single-scale modeling and filtering. For this analysis, we fit models for latent state dimensions *n*_*x*_ ∈ {2, 5 : 5 : 20}. To select *n*_*x*_ for a given fold and a given signal combination, we divide the training data for that fold into an inner training set consisting of 80% of the training data and inner test set consisting of the remaining 20% of the training data. We then learn the model parameters using the inner training set and use them to predict the behavior in the inner test set. We then choose the *n*_*x*_ that results in the best behavior prediction accuracy on the inner test set.

### 4.5. Statistical analysis

All the statistical analyses for paired samples are done one-sided with Wilcoxon signed rank test. Significance is declared if the *P <* 0.05. In cases where multiple comparison are being made, we use the false discovery rate (FDR) control procedure from Benjamini–Hochberg [122] to correct for all comparisons and report the FDR-corrected *P* values.

## Author contributions

P.A. and M.M.S. conceived the study and developed the new learning algorithm with help from O.G.S. P.A. performed all analyses. P.A., O.G.S, and M.M.S. wrote the manuscript. B.P. designed and performed the experiments for the nonhuman primate dataset. M.M.S. supervised the work.

## Acknowledgments

The authors acknowledge support of the Army Research Office (ARO) contract W911NF1810434 under the Bilateral Academic Research Initiative (BARI), NSF CAREER Award CCF-1453868, NIH DP2-MH126378, and NIH R01MH123770. We would like to thank Yuxiao Yang and Hamidreza Abbaspourazad in the Shanechi Lab for the valuable initial discussions.

## Appendices

## Appendix A. Transformation of moments for estimating

Σ^*zy*^. Our goal is to compute Σ^*zy*^ = Cov[**z** ^*f p*^, **y** ^*f p*^] (section 4.2.3) in terms of first and second moments of spiking and field potential observation {Σ^*Ny*^, Σ^*NN*^, Σ^*yy*^, *μ*^*N*^, *μ*^*y*^}, which are directly computable from data.

We have:

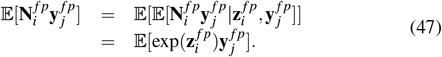

where ._*i*_ and. _*j*_ denote the *i*th and *j*th element of 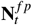 and 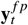, defined in section 4.2.3. Given that 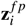 and 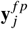 are jointly normal, i.e.:

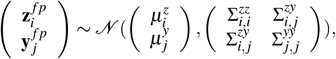

we can compute 𝔼 [exp(**z**_*i*_)**y** _*j*_] in equation 47 as:

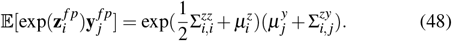

by taking a double integral required to compute 𝔼 [exp(**z**_*i*_)**y** _*j*_] with respect to the above bivariate normal distribution.

Further, we have:

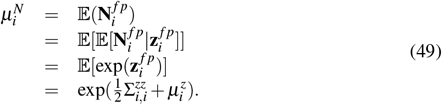

According to equations 47, 48 and 49, we have:

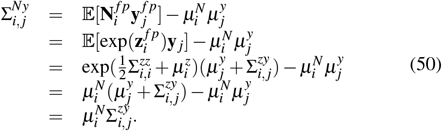

We then solve for 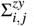 in equation 50, which gives:

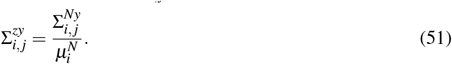

Of note, interestingly, the final relation in equation 51 resembles what reference [63] derives in their supplemental equation 6 as the moment transformation for relating Poisson observation to Gaussian *inputs*, as opposed to relating them to simultaneous Gaussian *observations*, which is derived here.

## Appendix B Constructing the future-past cross-covariance matrix H_*w*_ and auto-covariance matrix Λ_0_.

Having estimated 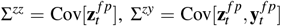 and 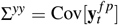 in section 4.2.3, we estimate **H**_*w*_ and Λ_0_ from them by definition, as:

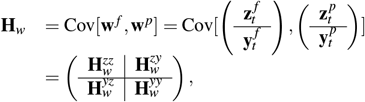

with

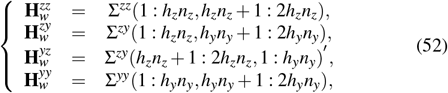

and

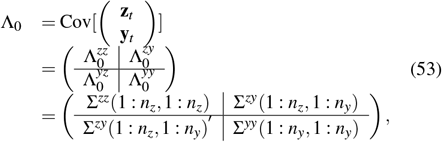

where : is the standard MATLAB operation for selecting some or all of rows and columns.

## Appendix C **Derivation of** 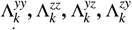 in equation 20 in terms of model parameters.

Given that noises **q**_*t*_ and **r**_*t*_ are white (equations 2-3), we know that **q**_*t*+*i*_ and **r**_*t*+*i*_ are orthogonal to **y**_*t*_ and **z**_*t*_ for all all positive integer *i*’s. Thus, we have:

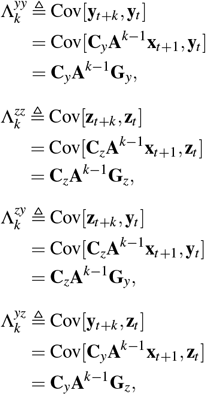

## Appendix D Derivation of the relation of noise covariance parameters to other model parameters.

Based on the multiscale dynamical model described in equation set 3, we can write the following equations for Cov[**x**_*t*_],

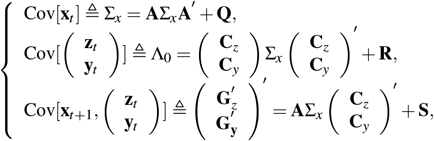

where 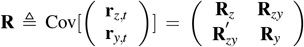 and 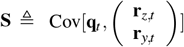. The relations in equation 40 can be easily obtained by rearranging the terms in the above equation set:

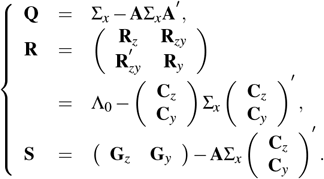

## Appendix E. Computation of Σ_*x*_ from model parameters.

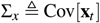 specifies the covariance of the latent state **x**_*t*_. Given the state transition matrix **A** and the state noise **Q** in equation 2, Σ_*x*_ can be computed analytically by solving the discrete Lyapunov equation:

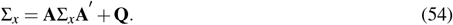

We use the “dlyap” MATLAB command to solve this equation and find Σ_*x*_.

## Appendix F. Generating C_*y*_ and C_*z*_ for simulating neural activity in section 4.3.

We generate 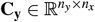 and 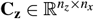 according to the following steps:

i. We first set the desired contribution of modes as follows:
  a. If mode *i* is a distinct spiking mode, i.e., only present in spiking activity (and absent in field potential activity), we set cntrb_*z,i*_ = 1 and cntrb_*y,i*_ = 0.
  b. If mode *i* is a distinct field mode, i.e., only present in field potential activity (and absent in spiking activity), we set cntrb_*z,i*_ = 0 and cntrb_*y,i*_ = 1.
  c. If mode *i* is a shared mode, i.e., present in both modalities, we set cntrb_*z,i*_ = 1 and cntrb_*y,i*_ = 1.
ii. We randomly generate entries of two matrices 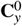 and 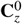 with elements uniformly distributed in the range [1, 1], whose dimensions are the same as **C**_*y*_ and **C**_*z*_, respectively.
iii. To adjust the required contribution of each mode in each modality (spiking or field potential activity), i.e., cntrb_*z,i*_ and cntrb_*y,i*_, we scale the columns corresponding to each mode in 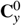 and 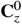, accordingly. We denote the scaled matrices as 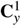 and 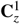. This leads to columns that correspond to distinct spiking (field) modes in 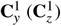 being filled with 0.
iv. To adjust the desired maximum firing rate for each neuron, we scale each row of 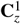, such that the upper bound of the analytical 95% confidence interval of the log firing rate, i.e., diag 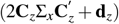, matches the log of desired maximum firing rates and denote it by 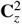.
v. Since contribution of each mode in the spiking activity may change after the previous step which scales the rows of 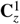, we recalculate the normalized contribution of each mode in the spiking activity nCntrb_*z,i*_ from 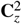. If the recomputed nCntrb_*z,i*_ is within an acceptable range of the original required contribution of each mode, we set 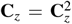 and go to the next step; otherwise, we go back to step (b) to regenerate 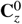. We set the acceptable range as [0.8, 1.2] times the original required normalized contribution.
vi. Finally, we scale the whole 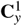 such that the sum of contribution of all the modes in the field potential activity is 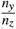 times that of the spiking activity (log firing rates) and denote it by **C**_*y*_. This step makes sure that the variance of log firing rates and field potential activity that is generated by the dynamical modes is proportional to the number of signals from each modality and the total variance is not dominated by one modality.

## Appendix G Estimating the similarity transform in section 4.3.2

The multiplication of the latent states by any invertible matrix **F**, also known as a similarity transform, gives an equivalent model with the same neural activity **y**_*t*_ and **N**_*t*_ but different basis for the latent state. More precisely, the set of parameters 𝒫 = {**A, C**_*z*_, **C**_*y*_, **G**_*z*_, **G**_*y*_, Λ_0_, **d**_*z*_, Σ_*x*_} with the latent state **x**_*t*_ will describe the same observations as a transformed model with latent state 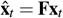 and parameters:

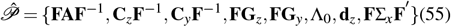

Thus, to evaluate the learned parameters against the real parameters for simulated models, we need to account for all the equivalent formulations for that simulated model. We address this with the same approach as used in our prior work [23].

To assess whether the learned parameters are close to any equivalent formulation of the true model or not, we first find the similarity transform 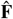 that makes the basis of the latent states for the identified model as close as possible to the basis of the latent states for the true model [23]. To do so, we first generate *T* = 10000 × *n*_*x*_ samples of neural activity from the true model. We then extract the predicted latent states using the multiscale filter [43] for both the true and identified models, denoted by 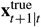 and 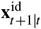, respectively. We then find the similarity transform **F** by minimizing the mean-squared error between the true predicted latent states and the transformed identified latent states, i.e.,

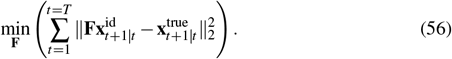

The analytical solution of this least squares problem is 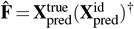, where 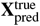 and 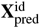 are matrices whose *t*th column contains 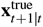 and 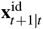, respectively. Having computed 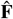, we apply it to the identified model parameters (equation 55) to get an equivalent model in a basis closer to that of the true model. We can then compare the true and the transformed identified model parameters (section 4.3.2).

## Appendix H Computing the error of learning the eigenvalues of A (i.e., normalized mode error) in simulations.

To compare the eigenvalues of the identified and true **A** matrices by computing the normalized error measure (equation 43), we first need to find a consistent ordering for the vectors containing the learned and true eigenvalues. This is because all reorderings of the eigenvalues can be thought of as diagonal elements of **A** in different *equivalent* models in canonical form (i.e., with block-diagonal **A** matrices [117]) that can be obtained with similarity transforms from each other (section 4.3.2). To match the ordering of the true and learned eigenvalue vectors, as in prior work [23], among all the possible *n*_*x*_! permutations of eigenvalue vector of **A**^id^, we find the closet one to the eigenvalue vector of **A**^true^ in terms of Euclidean distance. We then use this vector and compare it with the eigenvalue vector of **A**^true^ by computing the normalized error measure (equation 43).

**Figure A1.**
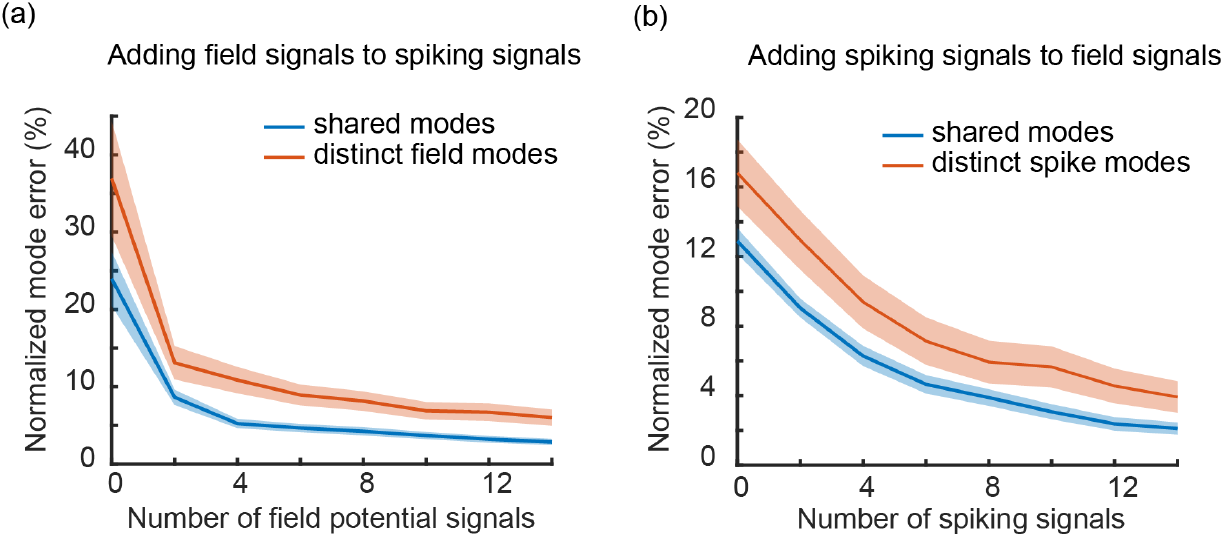
Multiscale SID (compared to single-scale SID) improves the identification of collective dynamics in both modalities. (**a**) Normalized errors of shared modes (blue) and distinct field modes (orange) that are only present in the field potential activity, as field potential signals are added gradually to 6 spiking signals. The start of the curves (i.e. 0 on x-axis) indicate normalized mode error for single-modal signals (i.e. spiking signals only). Solid line indicates mean across 50 simulated neural network activities and shaded area represents s.e.m. (**b**) same as (a) but for adding spiking signals to field potential signals. These simulation results suggest that multiscale SID correctly combines information across modalities of neural activity.

**Figure A2.**
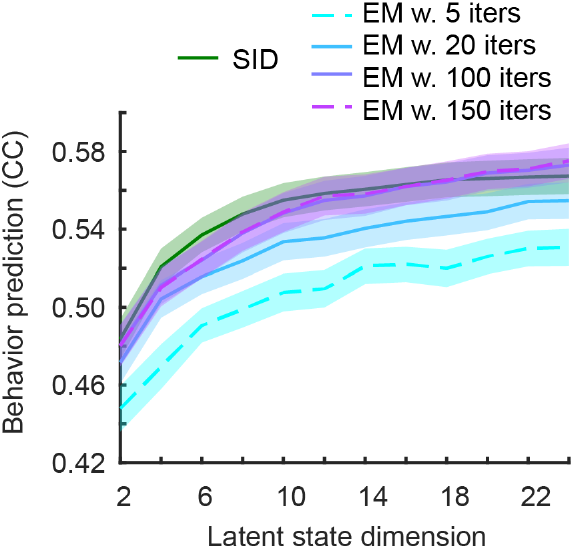
While being more computationally efficient and achieving better spike and field potential prediction (Figure 6), multiscale SID also matches the accuracy of converged multiscale EM in prediction of behavior in the NHP dataset. Behavior prediction accuracy quantified by correlation coefficient (CC) between the predicted and true behavior as a function of latent state dimension using multiscale SID and multiscale EM with different number of allowed iterations.

